# Rhomboid protease Rhbdl2 influences macrophage recruitment and wound repair in zebrafish

**DOI:** 10.64898/2026.02.13.705804

**Authors:** Saroj Gourkanti, Gayathri Ramakrishnan, Jacqueline Cheung, Taylor J. Schoen, Yazmin Munoz, Rosa M. Chavez, Jan Dohnálek, Katie Martin, Matthew Lovett-Barron, Thomas Whisenant, Kvido Strisovsky, Sonya E. Neal

## Abstract

Wound repair requires tight control of immune cell behavior, yet the mechanisms that restrain immune-driven wound repair responses remain poorly defined. Here, we demonstrate that rhomboid intramembrane serine protease Rhbdl2 influences wound repair in zebrafish. We generated *rhbdl2* mutants using CRISPR-Cas9 and found that, although Rhbdl2 is dispensable for normal development, its loss triggers enhanced wound repair following injury. This regenerative phenotype is accompanied by increased macrophage migration speed and accumulation at the wound site, as well as elevated early apoptosis and cell proliferation. Proteomic analyses reveal increased Rac2 protein levels in *rhbdl2* mutants, which was previously identified as a regulator of leukocyte motility. Functionally, Rac2 morpholino–mediated knockdown in *rhbdl2* mutant larvae suppresses the elevated macrophage recruitment and enhanced tissue repair phenotype. Together, these findings identify Rhbdl2 as a modulator of macrophage recruitment to the wound site during tissue repair, with implications for inflammatory disease, fibrosis, and tumor–immune interactions.

## Introduction

Millions of people worldwide will suffer from chronic, nonhealing wounds over the course of their lifetime, representing a major and growing clinical burden (Sen et al., 2009). Effective wound repair requires the precise integration of spatially and temporally coordinated signaling events that operate at both the single-cell and multicellular levels, encompassing epithelial cells, immune populations, and the extracellular matrix (Gao et al., 2024; Rousselle et al., 2019; Sorg et al., 2017). These processes are orchestrated by tightly regulated ligand– receptor signaling networks that control inflammation, cell migration, proliferation, and tissue remodeling (Liu and Fang, 2025; Mendonce et al., 2025). While wound repair shares components with homeostatic developmental pathways, the processes of wound repair also possess unique regulators that function in different stages (Khan et al., 2018). The innate immune system plays an important role in modulating the wound repair process by orchestrating the recruitment and behavior of leukocytes, including macrophages and neutrophils (De Oliveira et al., 2016; Phillips et al., 2020). These cells work by balancing pro- and anti-inflammatory signaling at the wound. Following injury, stress cues initiated by neighboring keratinocytes rapidly recruit neutrophils, which amplify and propagate inflammatory signaling throughout the wound site (Wilgus et al., 2013; Xu et al., 2024; Zhu et al., 2021). These early events subsequently promote the recruitment of macrophages, which function to combat infection and drive a pro-reparative signaling program (Krzyszczyk et al., 2018; Minutti et al., 2017). Dysregulated leukocyte recruitment or aberrant immune cell behavior can impair wound resolution, contributing to chronic wounds and increased susceptibility to infection (Miskolci et al., 2019; Schilrreff and Alexiev, 2022). While cell-intrinsic regulators such as Rac family GTPases are essential for leukocyte migration and function (Deng et al., 2011; Mishra et al., 2023; Rosowski et al., 2016), emerging studies continue to reveal novel regulators that modulate extrinsic recruitment signaling during tissue repair (Golenberg et al., 2020; Houseright et al., 2020; Karnam et al., 2023; Mercado Soto et al., 2025, Preprint; Tsourouktsoglou et al., 2020). The intramembrane serine protease Rhbdl2 has recently emerged as an attractive candidate implicated in wound repair (Cheng et al., 2011; Johnson et al., 2017). It is a member of the highly conserved rhomboid superfamily (Adrain et al., 2011; Freeman, 2014; Strisovsky, 2013). The rhomboid superfamily comprises intramembrane proteases, which function in many biological processes such as membrane protein homeostasis (Began et al., 2020; Bhaduri et al., 2025; Fleig et al., 2012; Grieve et al., 2021), intercellular signaling (Lemieux et al., 2007; Stevenson et al., 2007; Urban et al., 2001), parasitic invasion (Baker et al., 2006; Brossier et al., 2005; Gandhi et al., 2020), and protein trafficking (Knopf et al., 2024; Paschkowsky et al., 2016; Wunderle et al., 2016). Rhomboid intramembrane proteases cleave substrates within the lipid bilayer through a conserved serine-histidine catalytic dyad (Ha et al., 2013; Tichá et al., 2018). While the overall structural architecture of rhomboid proteases is highly conserved, individual family members exhibit remarkable substrate specificity, targeting and cleaving distinct and nonoverlapping repertoires of substrates (Bhaduri et al., 2022; Gourkanti et al., 2025). Notably, Rhbdl2 localizes to the cell surface and is enriched along the plasma membrane (Lohi et al., 2004), positioning it ideally to regulate ligand availability and receptor activation in response to tissue injury (Noy et al., 2016; Pascall and Brown, 2004; Zettl et al., 2011). Current knowledge of Rhbdl2 function is largely based upon the identification of its substrate repertoire via proteomic approaches, which have shown that Rhbdl2 cleaves critical factors of wound repair such as thrombomodulin, DDR1, and Spint1 (Johnson et al., 2017). Moreover, in vitro wound assays in Rhbdl2 mutant HaCaT cells show a significant delay in wound repair (Cheng et al., 2011). Importantly, previous characterization of Rhbdl2 function thus far relied heavily on in vitro cell culture studies, and no vertebrate knockout model has been established. Zebrafish have the remarkable ability to repair various tissues such as heart, spinal cord, and skin, and these processes are conserved in humans (Arjmand et al., 2020; Cigliola et al., 2020; Greenspan et al., 2024). Furthermore, there are well-established wound repair assays in zebrafish along with genetic tools and established transgenic lines to allow for an extensive exploration of Rhbdl2 function on cellular behavior in a reparative context (Miskolci et al., 2019). Therefore, zebrafish is a well-suited vertebrate model for robust systematic analysis of Rhbdl2. To that end, we established that zebrafish possess a single, highly conserved ortholog of Rhbdl2. Using an in vitro cleavage assay in HaCaT cells, we demonstrate that zebrafish Rhbdl2 retains enzymatic activity comparable with its human counterpart, confirming functional conservation across species. Accordingly, we generated a *rhbdl2* mutant zebrafish line and used wound assays, fluorescent imaging, and proteomic profiling to characterize its role in vertebrate wound repair. While loss of Rhbdl2 does not perturb embryonic development or adult viability, *rhbdl2* mutants exhibit a striking phenotype during tissue repair, characterized by enhanced wound regeneration following injury. This mutant phenotype is recapitulated by specific chemical inhibition of Rhbdl2 with a ketoamide inhibitor in WT zebrafish, confirming specificity. To uncover the molecular basis of this response, we performed unbiased global proteomic profiling and identified a significant posttranscriptional increase in Rac2 protein levels in *rhbdl2* mutants, a key regulator of leukocyte dynamics. Live imaging and quantitative analyses in *rhbdl2* mutant larvae revealed robust accumulation of macrophages and, to a lesser extent, neutrophils at the wound site, actively driving the enhanced reparative response observed in *rhbdl2* mutants. Together, these findings identify Rhbdl2 as a previously unrecognized regulator of injury-induced tissue repair, acting through modulation of immune cell dynamics.

## Results

### Zebrafish rhbdl2 functions as a canonical rhomboid protease

To investigate the role of the rhomboid protease Rhbdl2 in zebrafish, we characterized the annotated zebrafish gene *rhbdl2*. Our analysis identified transcripts with 8 exons (205, 202, and 204), 6 exons (203), and 7 exons (201) (**Fig. S1 A**). The *rhbdl2* gene encodes a 294–amino acid protein (GenBank: AAI64054.1) that is highly conserved among vertebrates, with homologous sequences present in lower eukaryotic and prokaryotic organisms. Sequence alignment of human and zebrafish Rhbdl2 polypeptides from the canonical transcript 205 demonstrated a high level of identity (64%), where zebrafish Rhbdl2 possesses the conserved serine-histidine dyad essential for proteolytic activity. Additionally, key rhomboid motifs, including WR and GxxxG, which are conserved across all rhomboid proteins (Kandel and Neal, 2020; Lemberg and Freeman, 2007), are retained in zebrafish Rhbdl2 (**Fig. S1 B**). Structural modeling using AlphaFold demonstrated that zebrafish Rhbdl2 retains the same fold as the human Rhbdl2 homolog (**Fig. S1 C**). Additionally, in situ hybridization data from the ZFIN database, together with single-cell RNA-sequencing data from Zebrahub, indicate that *rhbdl2* is expressed in the zebrafish epidermis throughout development beginning at 2 days postfertilization (dpf) (data not shown). To assess the proteolytic activity of zebrafish Rhbdl2, we subcloned its open reading frame (ORF) into a pcDNA3.1 expression vector and transfected HEK293 cells with either human or zebrafish Rhbdl2. Expression of both human and zebrafish Rhbdl2 was readily detected in cell lysates. Using a cleavage-based assay, when cells are transfected with either human or zebrafish Rhbdl2, we observed the release of a cleavage product of the well-characterized human Rhbdl2 substrate Spint1 into the culture medium, whereas cells transfected with empty vector showed only full-length Spint1 retained in the lysate (**Fig. S1 D**). Recent work identified EGFR as an endogenous substrate of human Rhbdl2 in keratinocytes, impacting cell migration and keratinocyte development (Johnson et al., 2026). Similar to human Rhbdl2, zebrafish Rhbdl2 proteolytically released the ectodomain of endogenous EGFR from the human keratinocyte cell line HaCaT deficient in its endogenous human Rhbdl2 (**Fig. S1 E**). Based on these findings, we concluded that zebrafish *rhbdl2* functions as a canonical rhomboid protease, maintaining conserved activity similar to its mammalian counterpart.

**Figure S1.**
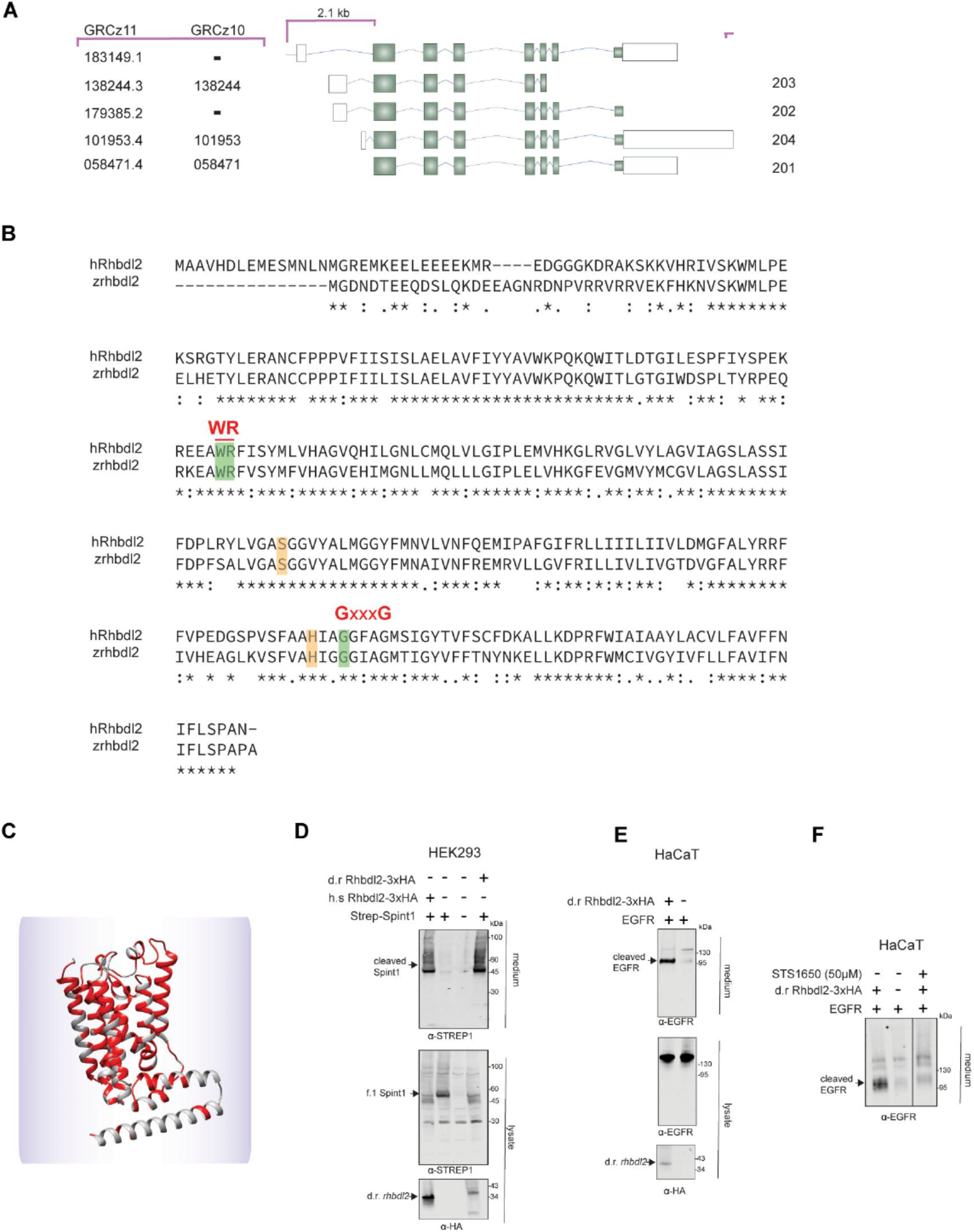
Characterization of zebrafish Rhbdl2 structure and function. (A) Diagrams of *rhbdl2* transcripts, where exons are displayed as solid boxes while introns are the connecting lines. Left: A list of ENSEMBL IDs with the last seven digits from both GRCz11 and GRCz10 genome assemblies (full accession IDs listed in Materials and methods). Right: List of transcript reference names from GRCz11. Asterisk indicates the canonical transcript discussed in this paper, while arrows highlight the sites of CRISPR-Cas9 guide mutagenesis, with the distance between them indicated. (B) Amino acid sequence comparison between human Rhbdl2 and zebrafish Rhbdl2 (transcript 205). Conserved rhomboid motifs (WR and GxxxG) are highlighted in green, and the serine-histidine catalytic dyad residues are highlighted in orange. (C) AlphaFold model depicting overlap between human and zebrafish Rhbdl2. Conserved residues are displayed in red. (D) Western blot of transiently transfected HEK293T cell culture conditioned media and lysate probed with Strep-(Spint1) (Qiagen, 34850, mouse, RRID: AB_ 2810987) and HA-(Rhbdl2) antibodies (Cell Signaling Technology, 3724 rabbit, RRID: AB_1549585). The top blot detects the Spint1 ectodomain shed by cells, while the middle and bottom blots display Spint1 and Rhbdl2 detection in lysates. N = 3 replicates. (E) Western blot of conditioned media and lysate from *rhbdl2*-deficient HaCaT cells stably expressing zebrafish Rhbdl2, probed with antibodies against endogenous human EGFR ectodomain (Abcam, ab264540, mouse, RRID: AB_3665828) and HA (Rhbdl2). The top blot detects shed EGFR ectodomain in conditioned media, while the middle and bottom blots display EGFR and Rhbdl2 in lysates. N = 3 replicates. (F) Western blot of conditioned media from *rhbdl2*-deficient HaCaT cells stably expressing zebrafish Rhbdl2 and treated with vehicle or the ketoamide inhibitor STS1650 (50 µM), probed with antibody against endogenous EGFR. STS1650 prevents zebrafish Rhbdl2-mediated EGFR ectodomain shedding. N = 3 replicates. Source data are available for this figure: SourceData FS1.

### An RNA-less rhbdl2−/− allele does not impact zebrafish development

To investigate the function of the *rhbdl2* gene, we generated an RNA-null allele using CRISPR-Cas9 to minimize compensatory mechanisms. Two gRNAs were designed to delete 2.1 kb of the upstream promoter region, exon 1, and the translation start site of exon 2, effectively preventing mRNA production (**Fig. 1 B**). The resulting *rhbdl2* heterozygous founders were identified by Sanger sequencing and outcrossed to generate stable heterozygote animals. These fish were subsequently inbred to obtain homozygous mutants (*rhbdl2*−/−). qPCR analysis using primers spanning exon 2–3 and exon 4–5 confirmed the absence of *rhbdl2* mRNA expression in zebrafish larvae at 2 and 6 dpf (**Fig. 1 C**). Because the rhomboid superfamily comprises multiple rhomboid proteases, we next sought to determine whether *rhbdl2*−/− zebrafish trigger redundancy or genetic compensation by other rhomboid family members. We quantified the mRNA levels of rhbdl1, rhbdl3, and rhbdl4 (**Fig. 1 D**). Notably, despite the genetic loss of *rhbdl2*, the expression levels of other rhomboid family members remained unchanged. Next, we examined whether *rhbdl2*−/− zebrafish exhibited any morphological abnormalities. Across different developmental stages, no visible defects were observed in 2 dpf larvae as well as in both adult females and males (**Fig. 1, E and F**). Collectively, these findings indicate that the loss of *rhbdl2* does not impact zebrafish development.

**Figure 1.**
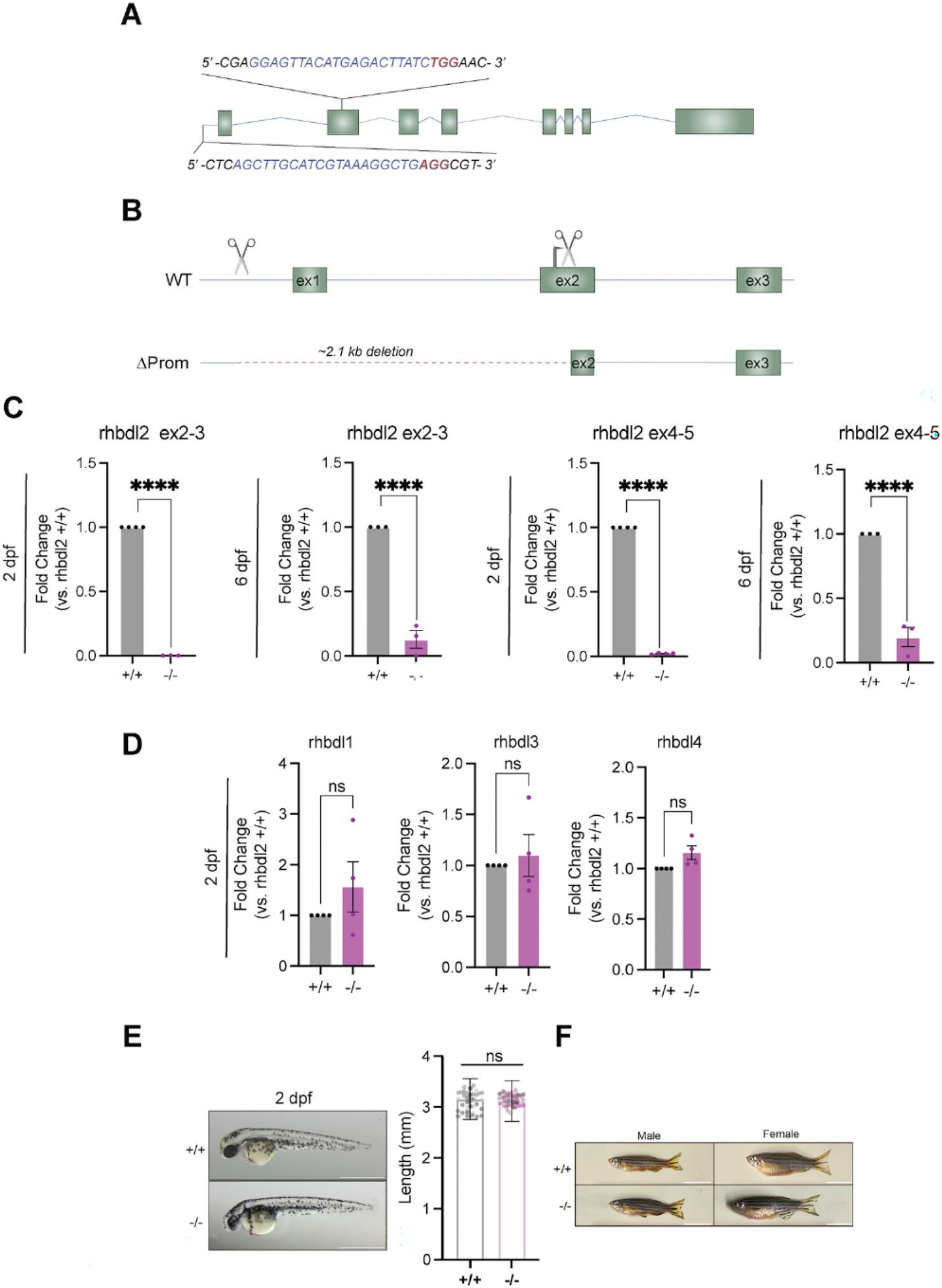
*rhbdl2* promoterless knockout zebrafish model does not display developmental defects. (A) Schematic of the *rhbdl2* gene with 5′ gRNA sequence and exon 2 gRNA sequence highlighted for CRISPR/Cas9 mutagenesis. The sequence of the gRNA is displayed in blue, and the protospacer adjacent motif is displayed in red. (B) Sequence comparison of WT and ΔProm knockout allele illustrating a 2.1 kb deletion. CRISPR/Cas9 guide sites are highlighted by scissors and translation start site is shown by a black arrow. (C) RT-qPCR of *rhbdl2* using primers spanning exon 2–3 and exon 4–5 at 2 dpf and 6 dpf larvae. (D) rhomboid family proteases rhbdl1, rhbdl3, and rhbdl4 in pooled homozygous cousin larvae from WT and ΔProm homozygous sibling incrosses. Data are from four pooled replicates with n = 10–20 larvae per sample. Means and SEM are reported, and a one-sample t test performed. ****, P < 0.0001. (E) Representative full-body bright-field images of 2 dpf larvae, WT cousin (top) and *rhbdl2*−/−(bottom). Scale bar = 1 mm. Quantification of full body length between WT and *rhbdl2*−/− larvae at 2 dpf. Data are from three independent replicates, with lsmeans (±SEM) reported and P values calculated by ANOVA with Tukey’s multiple comparisons, n = 43 (+/+), 46 (−/−). (F) Representative full-body images of 6 mpf adult zebrafish cousins with males (left) and females (right) indicated. Data from two independent replicates; scale bar = 10 mm.

### Loss of Rhbdl2 induces enhanced tail fin regeneration following injury

To examine the role of Rhbdl2 in wound repair, we performed tail transections on 3 dpf larvae at the tip of the notochord. Homozygous *rhbdl2* mutant and WT offspring were generated from incrosses of F3 or F4 adult WT or *rhbdl2*−/− siblings. This cousin-based breeding strategy, where homozygous WT and mutant adults are generated from the same heterozygous parents and incrossed separately, ensures that experimental WT and *rhbdl2*−/− larvae share equivalent genetic backgrounds while avoiding heterozygous siblings in experimental clutches. Both *rhbdl2*−/− and WT cousins underwent tail transection, and the wound repair area (measured from the vessel edge to the caudal edge of the tail fin) was evaluated beginning at 3 days post-wounding (dpw) (**Fig. 2 A**). To assess whether wounding itself alters *rhbdl2* transcript levels, we performed RT-qPCR on WT and *rhbdl2*−/− unwounded and 24 hpw larvae, which revealed no significant changes in expression (**Fig. 2 B**). Notably, *rhbdl2*−/− mutants displayed a significant increase in wound repair area compared with WT cousins at 72 hpw and 96 hpw (**Fig. 2 C**). To acutely inhibit Rhbdl2 activity and determine whether pharmacological inhibition recapitulates the mutant phenotype, we used the human Rhbdl2 inhibitor STS1650 (compound 11 from [Bach et al., 2024]), a peptidyl α-ketoamide that covalently and reversibly targets the active site serine residue of rhomboids (Bach et al., 2024; Johnson et al., 2026; Tichá et al., 2017). To first confirm that STS1650 effectively inhibits zebrafish Rhbdl2 proteolytic activity, we treated Rhbdl2-deficient HaCaT cells transfected with zebrafish Rhbdl2 with STS1650 (50 µM) or vehicle and found that STS1650 indeed abolished zebrafish Rhbdl2-mediated EGFR ectodomain shedding into conditioned media (**Fig. S1 F**). Having confirmed its efficacy against zebrafish Rhbdl2, we proceeded with the in vivo treatment of zebrafish larvae beginning at 2 dpf and performed tail transections at 3 dpf. WT larvae were treated with the Rhbdl2 inhibitor STS1650 at 10, 15, and 25 µM, and caudal fin fold regeneration was quantified at 72 hpw. Treatment with STS1650 increased regenerated fin fold area in a dose-dependent manner compared with DMSO-treated WT controls (**Fig. 2 D**). At 10 µM, STS1650 did not significantly increase regenerated area relative to +/+ DMSO controls. However, at 15 µM, a significant increase was observed, and this effect was further pronounced at 25 µM, where STS1650-treated WT larvae exhibited regenerated areas comparable with those of *rhbdl2*−/− larvae (**Fig. 2 D**). These results indicate that the enhanced wound repair phenotype is attributable to loss of Rhbdl2 proteolytic activity and is unlikely to be due to off-target effects of CRISPR-Cas9 mutagenesis. Given Rhbdl2’s role in caudal fin wound repair, we next examined whether it is required for fin development. Whole tail measurements of 6 dpf larvae that were age-matched to wounding experiments revealed no significant differences between WT and *rhbdl2*−/− mutant (**Fig. 2 E**). These findings suggest that Rhbdl2 is required for the regulation of caudal fin regenerative outgrowth but not for its development. To determine whether loss of Rhbdl2 alters large-scale tissue architecture during tissue repair, we quantified contour length as a proxy for collagen projections, a metric that reflects extracellular matrix organization and has been shown to correlate with effective wound repair capacity (LeBert et al., 2018; Miskolci et al., 2019). This analysis revealed no significant difference in contour length between WT and *rhbdl2*−/− mutants (**Fig. 2 F**). Given that fin contour length was indistinguishable between WT and *rhbdl2*−/− larvae, we next asked whether loss of Rhbdl2 affects more subtle aspects of ECM organization during wound repair. We utilized a cell-impermeable ECM-specific dye, Rhobo6, to visualize the collagen matrix in the repairing fin at 72 hpw (Fiore et al., 2025). Quantitative phenotypic scoring of collagen recovery did not reveal statistically significant differences among genotypes (**Fig. S2, A and B**). Together, these data indicate that loss of Rhbdl2 does not result in aberrant wound repair but instead is associated with enhanced wound repair.

**Figure 2.**
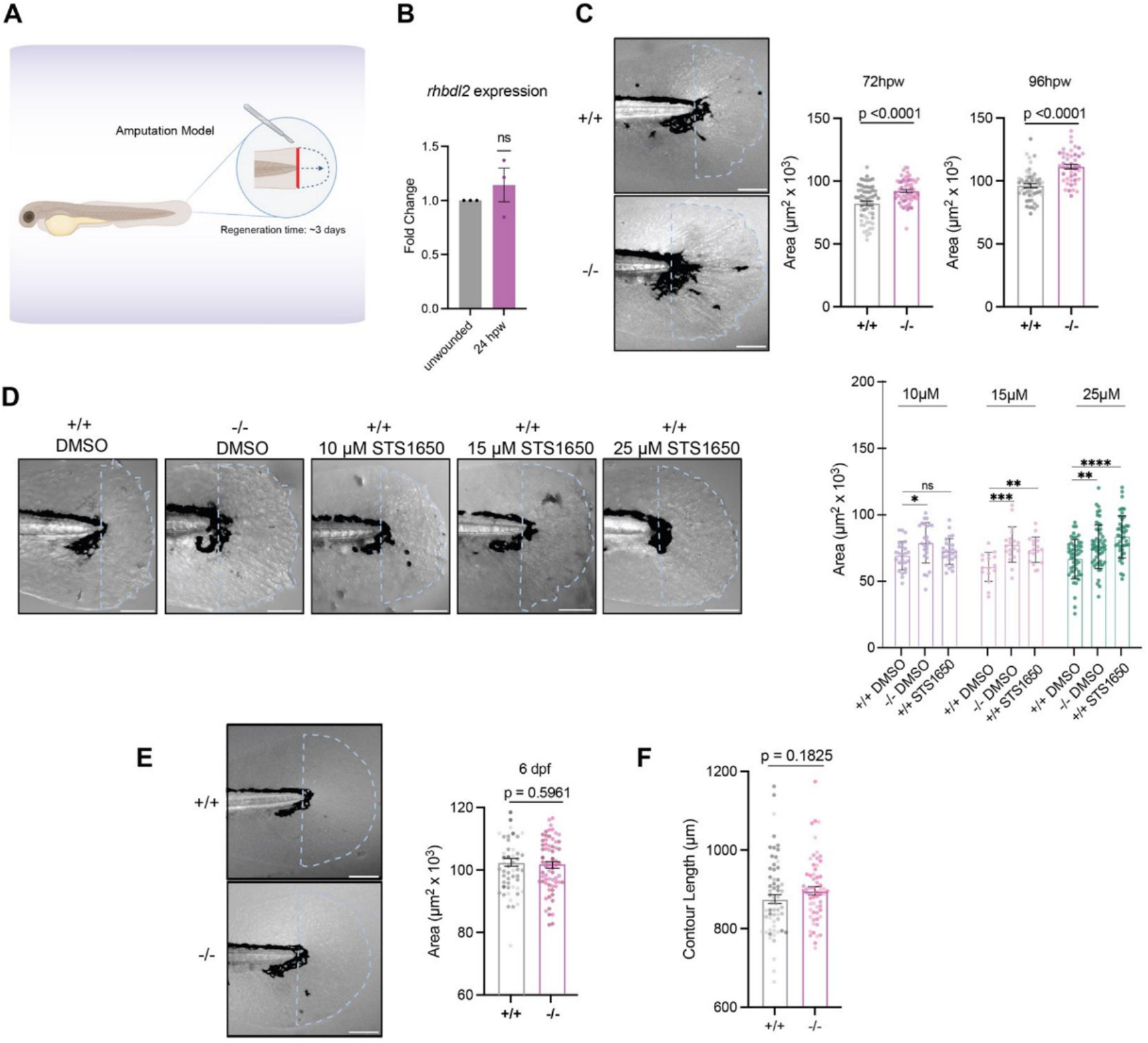
Loss of Rhbdl2 leads to enhanced wound repair. (A) Cartoon displaying transection wound assay conducted on 3 dpf larvae. Transections were performed at the tip of the notochord, and general wound repair time is considered complete at 3 dpw. Fin areas were measured from the tip of the notochord. (B) RT-qPCR of *rhbdl2* expression in *rhbdl2* (+/+) and *rhbdl2* (−/−) unwounded and 24 hpw larvae. Data are from three independent replicates, with lsmeans (±SEM) reported and a one-sample t test performed. No significant differences were detected between conditions. (C) Representative bright-field images of wound repair at 72 hpw and 96 hpw from three independent replicates, with lsmeans (±SEM) reported and P values calculated by ANOVA with Tukey’s multiple comparisons; 72 hpw: n = 59 (+/+), 61 (−/−), 96 hpw: n = 52 (+/+), 46 (−/−). (D) Pharmacological inhibition of *rhbdl2* phenocopies the *rhbdl2*− /− regeneration defect in a dose-dependent manner. Representative bright-field images of regenerating caudal fin folds at 72 hpw from WT (+/+) larvae treated with DMSO vehicle, *rhbdl2*−/− (−/−) larvae treated with DMSO, and WT larvae treated with 25 µM STS1650. Quantification of regenerated fin fold area (µm2 × 103) is shown on the right for each treatment group across concentrations. Data are from 3 independent replicates, with lsmeans (±SEM) reported; (+/+) DMSO: n = 26,16, 50, (−/−) DMSO: n = 25, 19, 53, (+/+) 10 µM: n = 26, (+/+) 15 µM: n = 17, (+/+) 25 µM: n = 39, and P values calculated by one-way ANOVA with Tukey’s multiple comparisons test. *P < 0.05, **P < 0.01, ***P < 0.001, and ****P < 0.0001; ns, not significant. Scale bar, 100 µm. (E) Representative bright-field images of unwounded tails age-matched at 6 dpf. Quantification of developmental fin area of 6 dpf larvae from three independent replicates with lsmeans (±SEM) reported and P values calculated by ANOVA with Tukey’s multiple comparisons, n = 52 (+/+), 68 (−/−). Scale bars = 100 µm. (F) Quantification of contour length in 72 hpw wounds in 2B. Contour length calculated as wound perimeter minus tail width at the notochord tip. Data are from three independent replicates with lsmeans (±SEM) reported, and P values calculated by ANOVA with Tukey’s multiple comparisons, n = 62 (+/+), 61 (−/−). Scale bars = 100 µm. hpw, h post-wounding.

**Figure S2.**
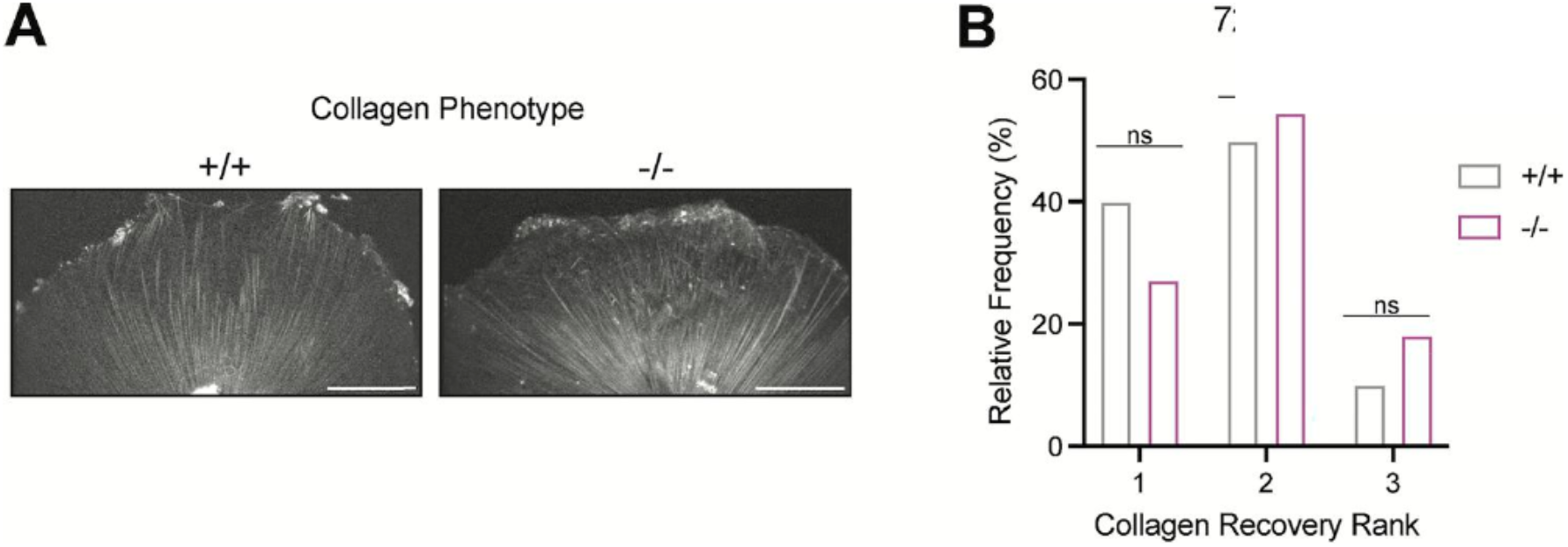
Characterization of the ECM matrix in wound-repaired fins of *rhbdl2*−/− mutants. (A) Representative images of WT or *rhbdl2*−/− wound-repaired fins at 72 hpw and labeled with Rhobo6. Images are average fluorescent z-stacks from two-photon microscopy, and levels were adjusted for best visualization. Scale bar, 100 µm. (B) Collagen scoring using a blinded phenotypic screening system across both genotypes as follows: 1 = little to no aberrations, 2 = moderate irregularities and scarring in collagen matrix, 3 = large irregularities or missing areas of signal. Data are from three independent experiments, n = 10 (+/+), 11 (−/−). Two-way ANOVA was conducted (P = 0.6667). hpw, h post-wounding.

### Rhbdl2-deficient larvae have increased wound-induced proliferation and apoptosis

Wound repair is a dynamic and coordinated process where the balance between cell proliferation and apoptosis is crucial for restoring the correct tissue architecture following injury (Aragona et al., 2017; Justynski et al., 2023; Zanca et al., 2022). To assess proliferative activity during fin wound repair, we performed 6-h EdU pulse labeling on 6 dpf larvae that were either left unwounded or subjected to tail transection and collected at 72 hpw. Representative merged bright-field and EdU (magenta) images are shown, with single EdU channels displayed in the right panel for clarity (**Fig. 3, A and C**). In both developmental (unwounded) and repairing fins, EdU-positive cells were detected predominantly within the distal fin fold region. Quantification of EdU-positive nuclei revealed no significant difference in proliferative index between WT and *rhbdl2*−/− larvae under unwounded conditions (**Fig. 3 B**). However, following injury, there was a robust increase in EdU incorporation throughout the repairing fin (**Fig. 3, D and E**), indicating elevated proliferative activity during tissue repair. To further quantify wound-induced proliferation, we focused on the dorsal region of the tail, as developmental proliferation is largely localized to the ventral fin (Golenberg et al., 2020). Within the dorsal region (white box in **Fig. 3 C**), *rhbdl2*−/− mutants exhibited a significant increase in proliferation following wounding compared with their WT cousins, whereas no differences were observed between unwounded WT and mutant fins (**Fig. 3 F**). These results suggest that Rhbdl2 specifically influences the proliferative response following injury. To complement our EdU incorporation analyses, we next assessed mitotic activity by co-staining wounded larvae with phospho-histone H3 (pH3, yellow), a marker for cells primed to undergo mitosis (**Fig. 3 G**). Consistent with our EdU findings, quantitative imaging revealed a significant increase in pH3-positive cells at 72 hpw in wound regions, but not at earlier time points (**Fig. 3 G**). To determine whether impaired wound repair in *rhbdl2*−/− larvae was also associated with altered cell death dynamics, we next examined apoptosis by staining for active caspase-3 at 24 and 72 h hpw (**Fig. 3 H**). Representative images show active caspase-3–positive signal (cyan) concentrated at higher levels during early wound repair stages. Quantification of the active caspase-3– positive area revealed a significant increase in apoptosis in *rhbdl2*−/− cousins compared with WT at both 24 and 72 hpw (**Fig. 3 H**). Importantly, this temporal pattern indicates that the increase in cell proliferation occurs subsequently to the onset of elevated apoptosis. Together, our findings indicate that loss of Rhbdl2 elicits an enhanced wound repair response characterized by increased cell proliferation and apoptosis.

**Figure 3.**
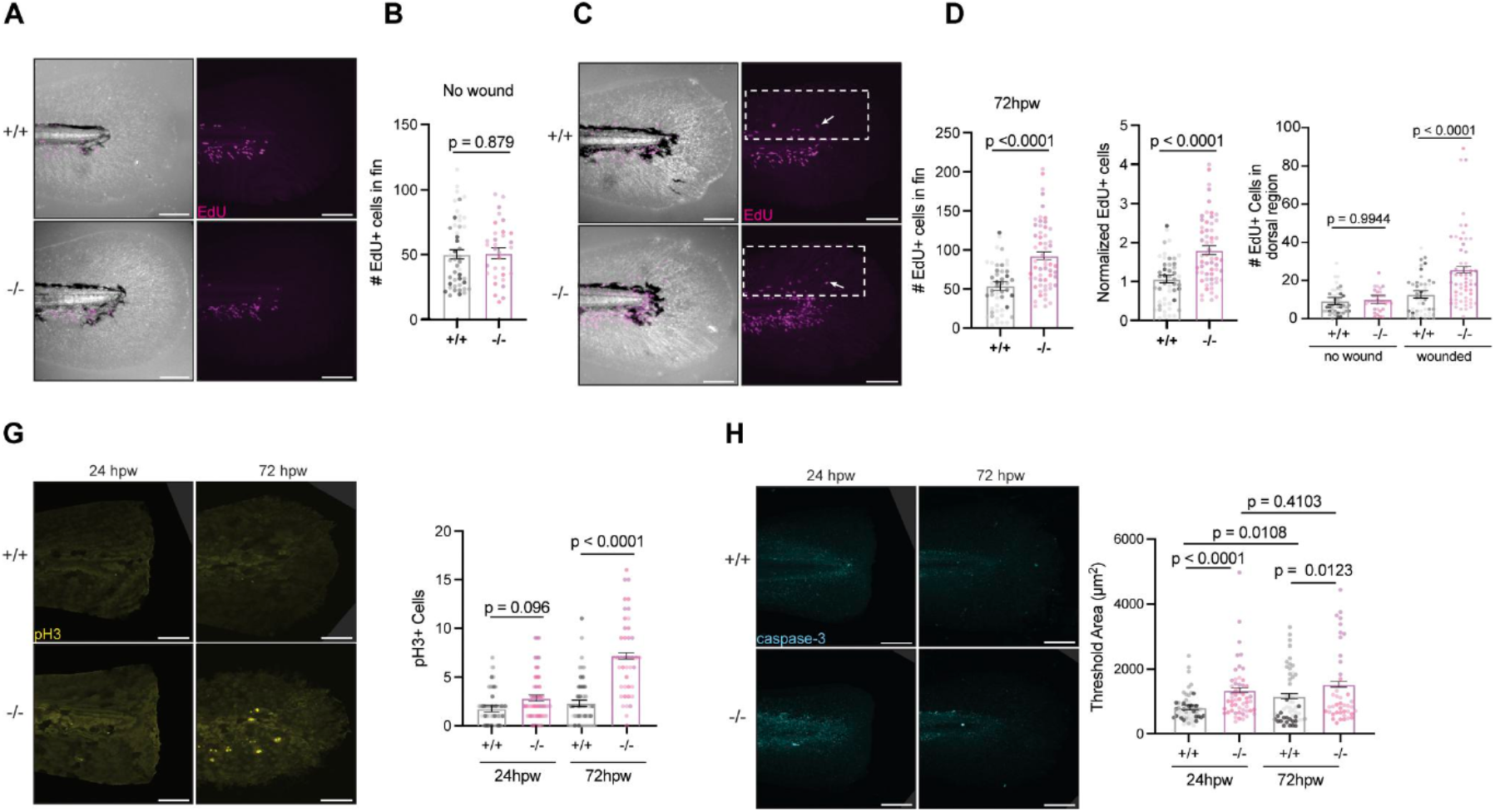
Loss of Rhbdl2 leads to increased proliferation and early apoptosis during wound repair. (A and C) Representative images of developmental unwounded (A) or 72 hpw (C) fins pulsed with EdU for 6 h. Merged bright-field and EdU (magenta) images are shown on the left, with a single EdU (magenta) image shown on the right. Dorsal region highlighted by a white box in C. (B and D) Quantification of EdU-positive cells in unwounded (B) and 72 hpw (D). (E) Number of EdU-positive cells in wounded fin normalized to respective unwounded conditions. (F) Quantification of EdU-positive cells within dorsal area of the fin. All EdU data are from four pooled independent replicates and cell counts were calculated from the region at tip of caudal blood loop through tail fin (no wound, n = 46 [+/+], 32 [−/−]; 72 hpw, n = 47 [+/+], 58 [−/−]). (G) Representative images of pH3 (yellow) stained tails at 24 hpw (left) and 72 hpw (right). Quantification of pH3-positive nuclei is shown on the right. (H) Representative images of active-Caspase3 (cyan) stained tails at 24 hpw (left) and 72 hpw (right). Quantification of active-caspase3 threshold area shown on the right. Data are from three independent experiments (24 hpw, n = 54 [+/+], 54 [−/−]; 72 hpw, n = 53 [+/+], n = 42 [−/−]). Lsmeans (±SEM) are reported, and P values calculated by ANOVA with Tukey’s multiple comparisons for all experiments. Scale bars = 100 µm. hpw, h post-wounding.

### Proteome profile of rhbdl2−/− zebrafish reveals alterations in Rac protein

Because Rhbdl2 functions as an intramembrane protease that directly cleaves and regulates the abundance of its substrates, we sought to determine how loss of Rhbdl2 impacts the global proteome in vivo. To obtain an unbiased, systems-level view of protein-level changes associated with Rhbdl2 deficiency, we performed label-free quantitative proteomics on zebrafish larvae. We focused our analysis at 2 dpf, a developmental stage at which Rhbdl2 expression is well established in epithelial tissues, based on in situ hybridization data from ZFIN.

We analyzed global proteome changes in 2 dpf zebrafish larvae, comparing WT (n = 150) and *rhbdl2*−/− mutant (n = 150) groups (**Fig. 4 A**). Multidimensional scaling analysis revealed that *rhbdl2*−/− mutant replicates clustered more closely together than WT (**Fig. 4 B**). However, a heatmap (**Fig. S3 A**), clustered based on protein abundance suggests a high degree of similarity in *rhbdl2*−/− mutant samples or WT larvae samples. Mass spectrometry identified a total of 8,932 Swiss-Prot protein id proteins, with 2,425 meeting significance thresholds for inclusion in statistical analysis. Among them, 48 proteins were differentially expressed, with 26 lower protein abundance and 22 exhibiting higher protein abundance in *rhbdl2*−/− mutants compared with WT (**Fig. 4 C** and **Fig. S4 A**). Notably, among the differentially expressed genes, bcam, cdh2, fras1, sccphdb, glg1a, glg1b, and plxnb2a encode factors involved in epithelial homeostasis and were also significantly enriched in a previous in vitro screen (**Fig. S3 B**) (Johnson et al., 2017). Gene ontology (GO) analysis indicated that the differentially regulated proteins were primarily associated with biological processes such as neutrophil and leukocyte migration, Rac protein signal transduction, apoptotic cell engulfment, and cell chemotaxis (**Fig. 4 D**). Rho GTPases are crucial regulators of innate immune cell functions, including migration and phagocytosis. The Rac subfamily of Rho GTPases consists of Rac1, Rac2, Rac3, and RhoG, each differing in expression patterns. Rac1 is ubiquitously expressed, Rac2 is primarily found in hematopoietic cell lineages, and Rac3 is predominantly expressed in neural tissue.

**Figure 4.**
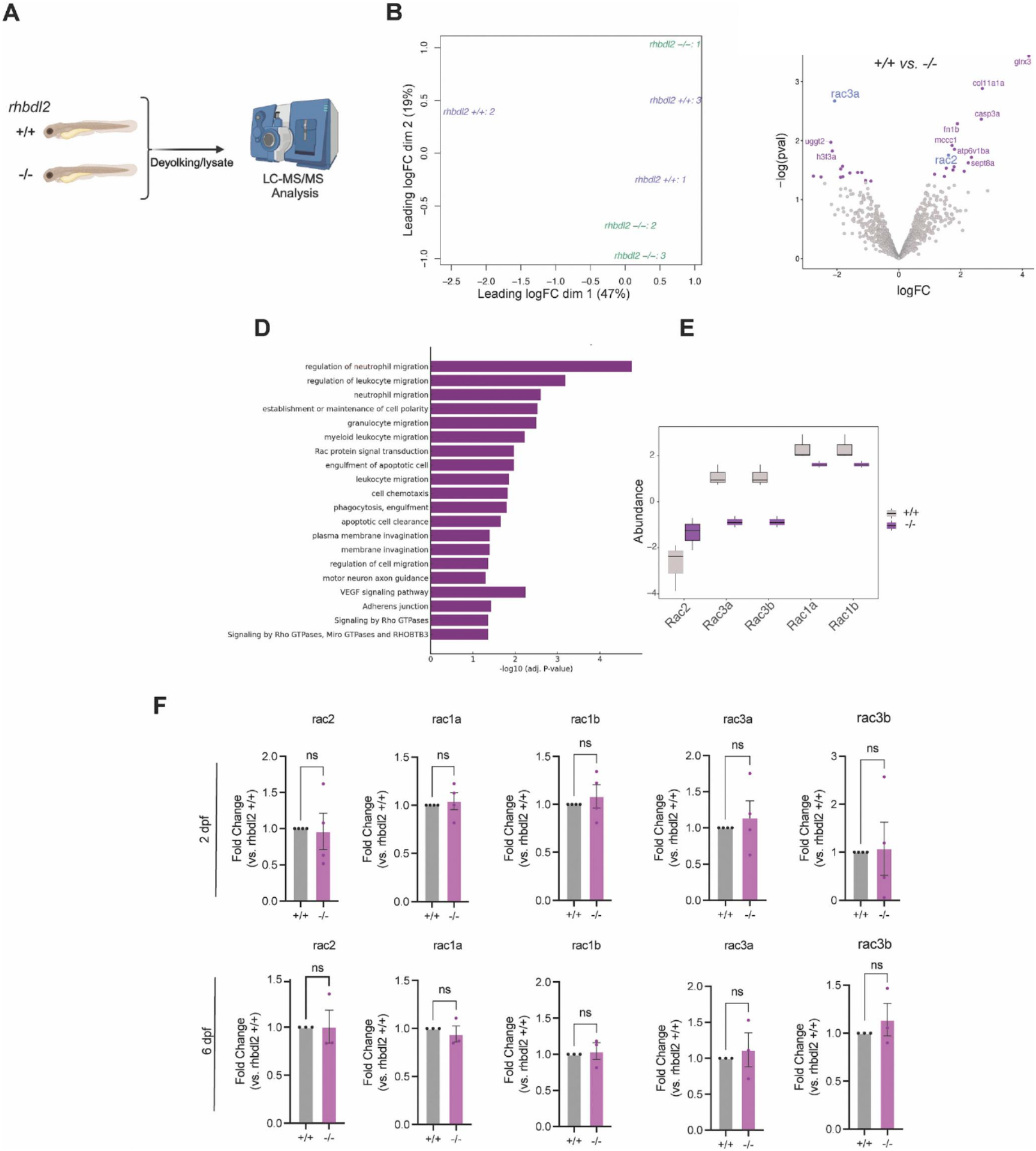
*rhbdl2*−/− mutants display differences in Rac protein levels. (A) Schematic depicting larval preparation of protein lysates for LC-MS/MS analysis. Data are from three independent experiments with n = ~50 larvae per sample (B) Multidimensional scaling (MDS) plot of the replicate WT (purple) and *rhbdl2*−/− (green) samples. The x-axis shows the separation of the samples on the first dimension and accounts for 47% of the variability in the data, while the y-axis shows the second dimension and accounts for 19% of the variability. (C) Volcano plot of the abundance of proteins in the limma comparison of WT and *rhbdl2*−/− samples. The x-axis shows the log fold change, and the y-axis represents the negative log of the comparison P value. Proteins with a significant P value (<0.05) are colored purple, and all others are colored white. The top 20 proteins (by P value) are labeled. (D) Pathway enrichment analysis using g:Profiler against cellular component GO terms. (E) Boxplot of the Rac proteins shows the abundance of each protein in the WT (white) and *rhbdl2*−/− (purple) samples. The box represents the interquartile range (IQR, 25–75%) with the median represented by a black line. The whiskers represent the range ±1.5× IQR. (F) RT-qPCR of Rac family genes on pooled homozygous cousin larvae from WT and ΔProm homozygous sibling incrosses. Data are from four pooled replicates with n = 10–20 larvae per sample. Means and SEM reported and unpaired t test performed.

**Figure S3.**
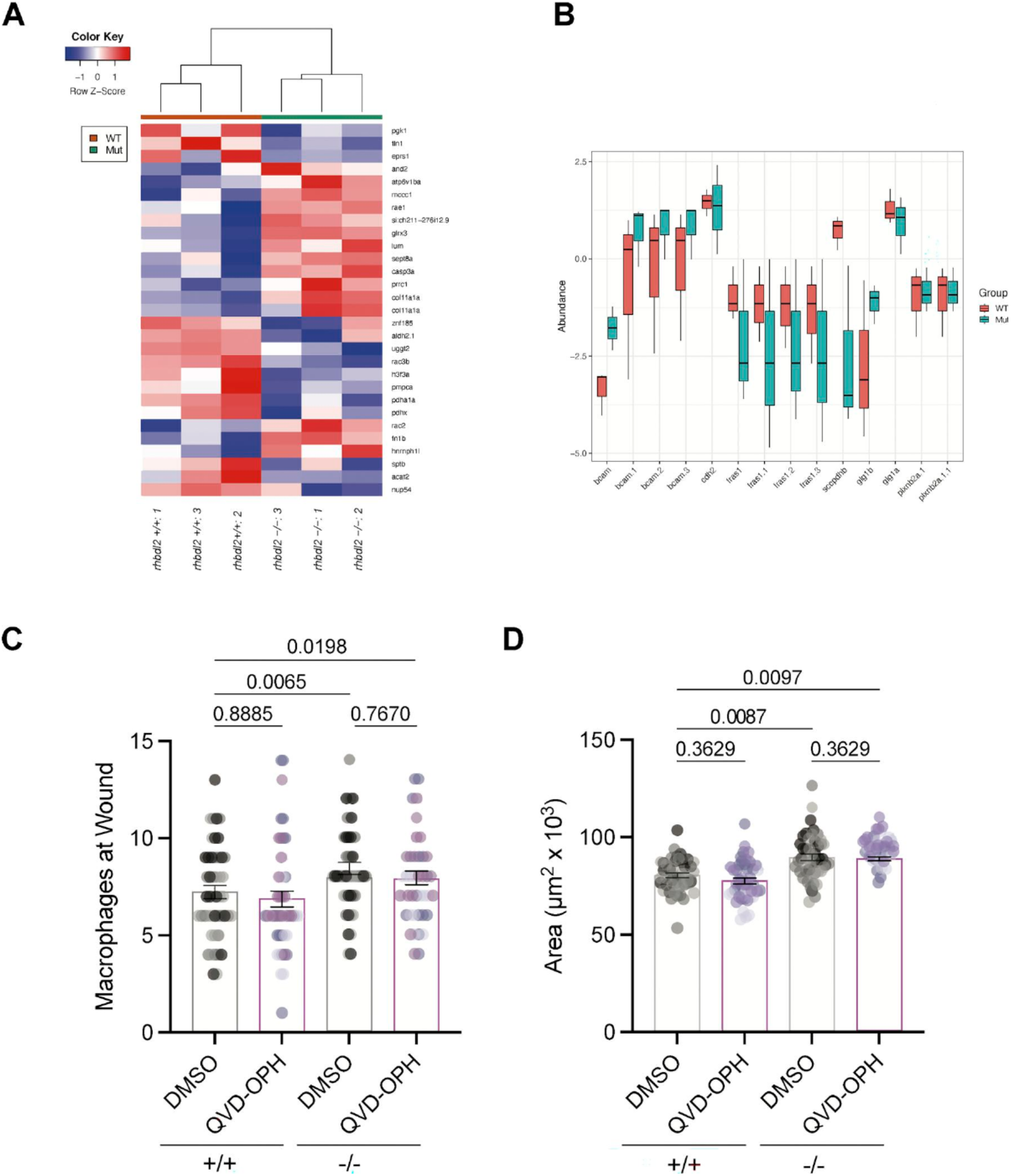
Differential abundance of proteins between WT and *rhbdl2*−/− larvae. (A) Heatmap of differentially abundant proteins that are detected in all WT and *rhbdl2*−/− samples. WT (orange) and *rhbdl2*−/− (teal) samples are represented across the top, along with a dendrogram showing the clustering of the samples based on the abundance of the proteins shown. (B) Boxplot of the proteins overlapping between the mass spectrometry proteomics results and Table 1 of (Johnson et al., 2017). For each protein, the proteomic abundance is shown for the WT (orange) and *rhbdl2*−/− samples (teal). The box represents the interquartile range (IQR, 25%–75%) with the median represented by a black line. The whiskers represent the range ±1.5× IQR. (C) Quantification of macrophage number at the wound site at 72 hpw in WT and *rhbdl2*−/−larvae treated with DMSO or the pan-caspase inhibitor QVD-OPH (50 µM). N = 3 independent replicates, n = 47 (WT DMSO), n = 48 (WT QVD-OPH), n = 47 (*rhbdl2*−/− DMSO), n = 44 (*rhbdl2*−/− QVD-OPH). Lsmeans (±SEM) reported; P values calculated by ANOVA with Tukey’s multiple comparisons. (D) Quantification of wound repair area at 72 hpw in WT and *rhbdl2*−/−larvae treated with DMSO or QVD-OPH (50 µM). N = 3 independent replicates, n = 47 (WT DMSO), n = 48 (WT QVD-OPH), n = 47 (*rhbdl2*−/− DMSO), and n = 44 (*rhbdl2*−/− QVD-OPH). Lsmeans (±SEM) are reported; P values calculated by ANOVA with Tukey’s multiple comparisons. hpw, h post-wounding.

**Figure S4.**
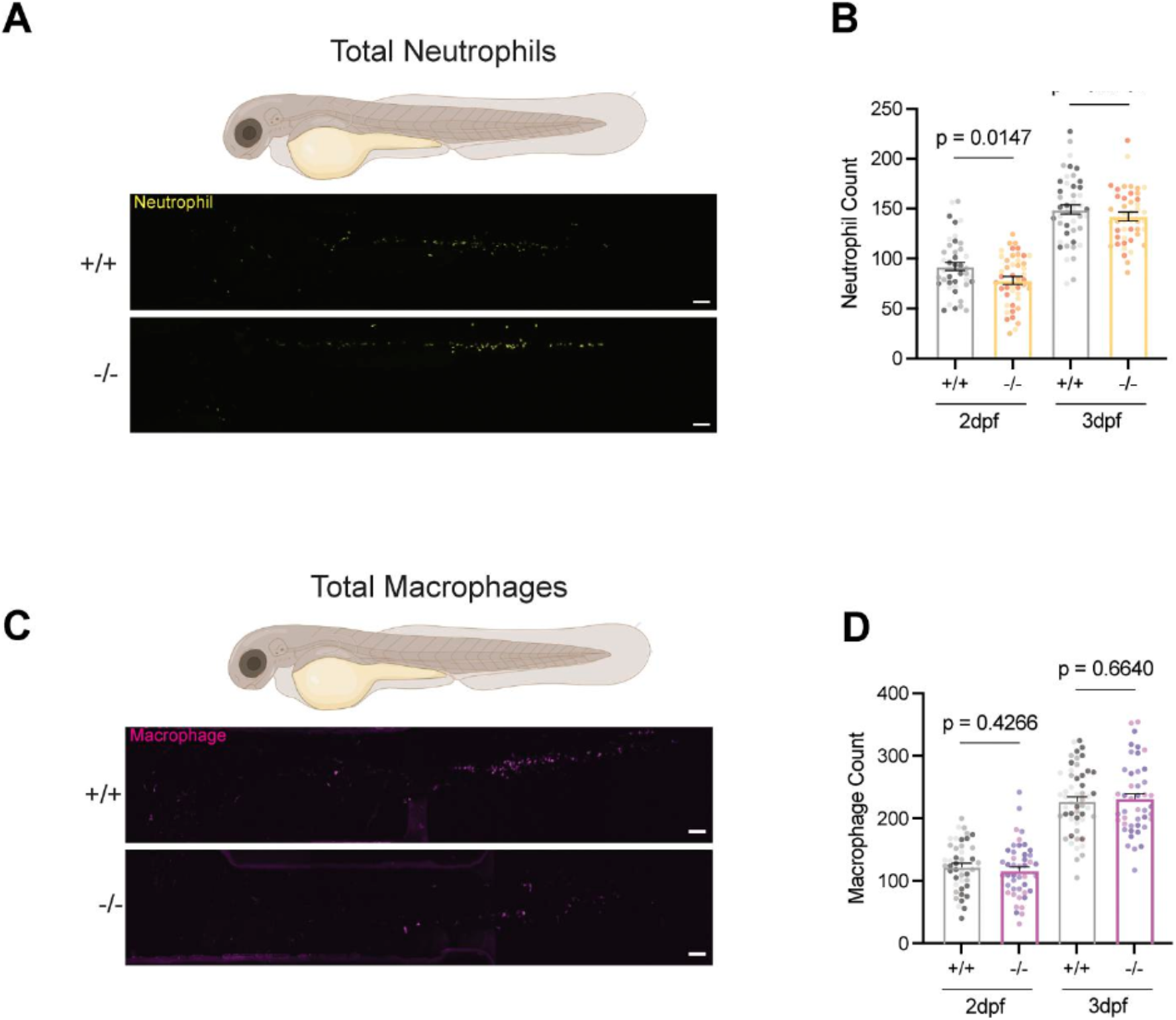
Development of leukocytes is unaffected in promoterless *rhbdl2*−/− mutants. (A) Representative full-body images of 2 dpf whole larvae with GFP-labeled neutrophils. (B) Quantification of the total number of neutrophils in whole larvae at 2 dpf and 3 dpf from three independent replicates (2 dpf, n = 43 [+/+], 46 [−/−]; 3 dpf, n = 45 [+/+], 45 [−/−]). (C) Representative full-body images of 2 dpf whole larvae with DsRed-labeled macrophages. (D) Quantification of total number of macrophages in whole larvae at 2 dpf and 3 dpf from three independent replicates (2 dpf, n = 44 [+/+], 44 [−/−]; 3 dpf, n = 52 (+/+), 44 [−/−]). Lsmeans (±SEM) are reported, and P values calculated by ANOVA with Tukey’s multiple comparisons for all experiments. Scale bar = 100 µm.

Rac2 constitutes 90% of the Rac protein in immune cells, where Rac1 and Rac2 share both redundant and distinct functions (Deng et al., 2011). Moreover, Rac proteins are well-characterized for playing a vital role in immune cell motility (Rosowski et al., 2016). Proteomics analysis using box plots confirmed that Rac1a, Rac1b, Rac3a, and Rac3b exhibited lower protein abundance, whereas Rac2 has higher protein abundance in *rhbdl2*−/− mutants compared with WT (**Fig. 4 E**). RT-qPCR analysis of Rac gene mRNA levels showed no significant changes, suggesting that alterations in Rac levels occur at the protein level rather than through transcriptional regulation (**Fig. 4 F**). Collectively, our proteomic analysis suggests that Rhbdl2 affects Rac GTPases at a posttranscriptional level, underscoring a potential role for Rhbdl2 in innate immunity.

### Loss of Rhbdl2 increases macrophage accumulation at wounds during tailfin wound repair

Our proteomic analyses suggest that Rhbdl2 modulates leukocyte migration, an important process contributing to overall wound recovery. Neutrophils are among the first immune cells to respond to epithelial injury, triggering inflammatory signals that promote the recruitment and proliferation of macrophages and other cell types necessary for pro-reparative wound repair. However, excessive neutrophil and macrophage accumulation or failure to migrate to the wound site can impair proper tissue repair (Miskolci et al., 2019). Accordingly, Tg(lyz:GFP; mpeg1:dsRed) reporter fish were crossed into the *rhbdl2*−/− mutant background to enable simultaneous visualization of neutrophil and macrophage recruitment to tail wounds during wound repair. We first focused on the initial recruitment of leukocytes to the wound using time-lapse microscopy from 1 to 6 hpw in the Tg(lyz:GFP; mpeg1:dsRed) background (**Fig. 5 A**). As expected, we observed a steady increase in both neutrophil and macrophage recruitment to the wound between 1 and 6 hpw (**Fig. 5 A** and Video 1). In *rhbdl2*−/− mutants, neither neutrophil nor macrophage recruitment was significantly impaired compared with WT. However, we observed a trend toward increased macrophage recruitment at 5–6 hpw, although this difference did not reach statistical significance (**Fig. 5, A–C** and Video 2). In addition, we assessed neutrophil and macrophage motility toward the wound by measuring their migration speed. For neutrophils, we detected no significant differences between genotypes, but interestingly, there was a modest increase in macrophage migration speed in *rhbdl2*−/− mutants in comparison with WT (**Fig. 5, D and E**). We next turned our attention to later stages of leukocyte recruitment. Transection wound assays were performed on transgenic cousins, and tails were fixed and imaged at 24, 48, and 72 hpw (**Fig. 5 F**). Neutrophil recruitment was comparable between WT and *rhbdl2*−/− larvae at 24 and 48 hpw, with only a modest elevation at 72 hpw in the mutant, in line with previous reports indicating that neutrophils primarily function during the early inflammatory stage of wounding (**Fig. 5 G**). In contrast to neutrophils, macrophage recruitment to the wound was markedly increased in *rhbdl2*−/− mutants at 24 and 48 hpw, followed by normalization by 72 hpw, indicating a prolonged but transient macrophage accumulation during the later stages of wound repair response (**Fig. 5, H and I**). We also observed elevated levels of the apoptotic factor Casp3a in the *rhbdl2*−/− proteome (**Fig. 4 C**), consistent with our finding of increased active caspase-3 staining in mutant fins during wound repair (**Fig. 3 H**). This raised the question of whether the enhanced macrophage recruitment and regenerative outgrowth in *rhbdl2*−/− mutants are secondary consequences of elevated apoptosis, for example, through efferocytotic macrophage recruitment driven by apoptotic cells, which in turn stimulates compensatory proliferation (Bhatt et al., 2022). To test this, WT and *rhbdl2*−/− mpeg:dsRed larvae tails were transected at 3 dpf, treated with the pan-caspase inhibitor QVD-OPH (Cambier et al., 2017; Caserta et al., 2003) (50 µM) or DMSO vehicle, and larvae were collected and fixed at 72 hpw. Quantification of macrophage accumulation at the wound site (**Fig. S3 C**) and regenerative area (**Fig. S3 D**) revealed that caspase inhibition did not suppress either macrophage recruitment or wound repair outgrowth in *rhbdl2*−/− mutants. These data suggest that macrophage accumulation at the wound site and the enhanced wound repair phenotype in *rhbdl2*−/− larvae are not simply downstream consequences of elevated apoptosis, and that Rhbdl2 influences wound repair through mechanisms that operate independently of caspase-mediated cell death. To determine whether these phenotypes reflected changes in leukocyte abundance, we quantified total neutrophils and macrophages in whole larvae. Neutrophil numbers were modestly reduced in *rhbdl2*−/− mutants at 2 dpf but were indistinguishable from WT by 3 dpf (**Fig. S4, A and B**). In contrast, total macrophage numbers did not differ between WT and *rhbdl2*−/− mutants at either 2 or 3 dpf (**Fig. S4, C and D**). Together, these data indicate that Rhbdl2 primarily modulates macrophage accumulation during the later phases of wound repair, rather than altering leukocyte development.

**Figure 5.**
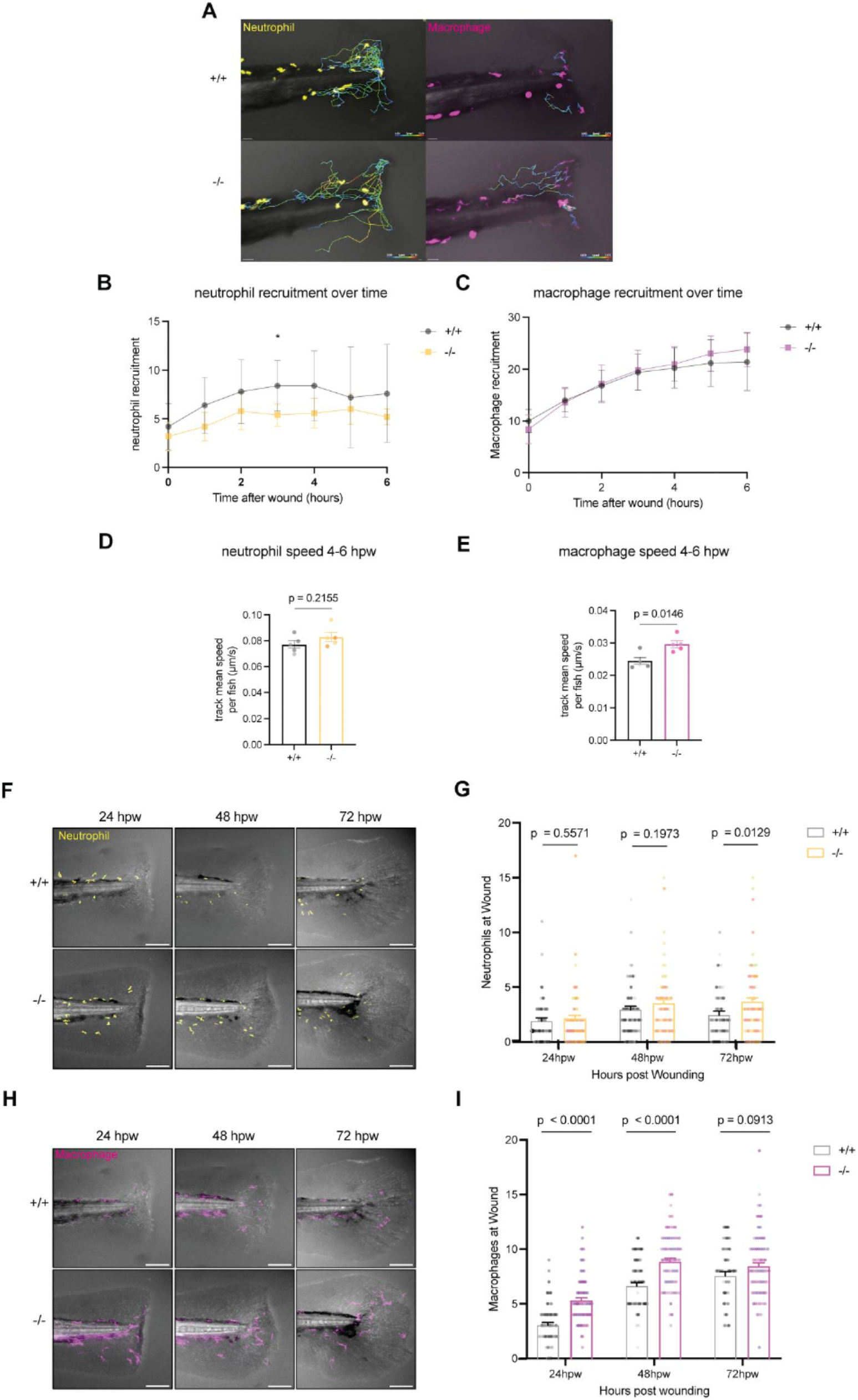
Loss of Rhbdl2 elicits increased retention of macrophages at the tailfin wound. (A) Representative serial images from 6 hpw time-lapse imaging comparing WT and *rhbdl2*−/− in (Tg(lyz:GFP;mpeg1:dsRed)) background; see Videos 1 and 2. Lines representing neutrophil (top) or macrophage (bottom) tracks over time are shown with warmer colors indicating faster speed. Data are from two independent replicates with n (larvae) = 5 (+/+), 5 (−/−), n (macrophages) = 43 (+/+), 46 (−/−), n (neutrophils) = 21 (+/+), 20 (−/−). (B and C) Quantification of the number of neutrophils (top) or macrophages (bottom) in the wound region distal to the caudal vein loop over the course of time-lapse imaging. (D and E) Quantification of instantaneous speed of neutrophils (D) and macrophages (E) during the later half (4–6 hpw) of time-lapse imaging; Lsmeans (±SEM) reported and P values calculated by ANOVA with Tukey’s multiple comparisons for all experiments. Scale bars = 30 µm. (F–H) Representative images of leukocytes at wounded tails from 24 to 72 hpw, visualized with GFP-labeled neutrophils (F) and DsRed-labeled macrophages (H) (Tg(lyz:GFP; mpeg1:dsRed)) merged with bright-field image. (G–I) Quantification of neutrophils and macrophages at the wound. All data from four independent replicates (24 hpw, n = 66 [+/+], 70 [−/−]; 48 hpw, n = 66 [+/+], 64 [−/−]; 72 hpw, n = 56 [+/+], 71 [−/−]). hpw, h post-wounding.

#### *Video 1. Go to* https://doi.org/10.17632/gwh3j3rrjr.2

*Time-lapse imaging of leukocyte recruitment to the tailfin wound in a WT larva. Confocal time-lapse of a WT larva Tg(lyz:GFP;mpeg1:dsRed imaged from 0 to 6 h post-wounding (hpw), showing neutrophils (yellow) and macrophages (magenta) migrating toward and accumulating at the tailfin transection wound. Corresponds to* ***Fig. 5 A***. *Scale bar, 30 µm*.

#### *Video 2. Go to* https://doi.org/10.17632/gwh3j3rrjr.2

*Time-lapse imaging of leukocyte recruitment to the tailfin wound in a rhbdl2−/− larva. Confocal time-lapse of a rhbdl2−/− larva Tg(lyz:GFP;mpeg1: dsRed imaged from 0 to 6 h post-wounding (hpw), showing neutrophils (yellow) and macrophages (magenta) migrating toward and accumulating at the tailfin transection wound. Corresponds to* ***Fig. 5 A***. *Scale bar, 30 µm*.

### Rac2 contributes to enhanced macrophage recruitment and wound repair in rhbdl2−/− larvae

Live cell tracking of macrophages in Rhbdl2-deficient larvae revealed a significant increase in macrophage recruitment to the wound site during later phases of wound repair. Proteomic analyses identified Rac2 as upregulated upon loss of Rhbdl2, suggesting that elevated Rac2 may contribute to these changes in macrophage recruitment. We therefore hypothesized that in *rhbdl2*−/− larvae, morpholino-mediated knockdown of Rac2 would restore macrophage recruitment to a WT-like state during wound repair. We employed a translation-blocking Rac2 morpholino previously validated in the literature, which has been shown to reduce both neutrophil and macrophage recruitment to the wound site (Deng et al., 2011). Indeed, Rac2 morpholino treatment in WT and *rhbdl2*−/− mutants significantly impaired neutrophil recruitment to the wound at all examined time points (24, 48, and 72 hpw) compared with control morpholino, confirming effective Rac2 knockdown (**Fig. 6, A and B**). Given that Rhbdl2 primarily influences macrophage recruitment during late-stage wound repair, we focused our analysis on macrophages. Consistent with earlier reports (Rosowski et al., 2016), depletion of Rac2 in WT larvae impaired macrophage recruitment to wounds (**Fig. 6, A and C**). As expected, at 24 and 48 hpw, *rhbdl2*−/− mutants injected with control morpholino exhibited a significant increase in macrophage recruitment at the wound compared with WT control. Strikingly, Rac2 knockdown in the *rhbdl2*−/− mutants significantly suppressed this phenotype, restoring macrophage recruitment to levels indistinguishable from WT (**Fig. 6, A and C**). At later time points (48 and 72 hpw), macrophage recruitment converged across genotypes and morpholino conditions. Together, these results suggest that the Rhbdl2-dependent effect of macrophage recruitment during wound repair is mediated through Rac2. We hypothesized that Rac2-dependent macrophage recruitment drives the enhanced reparative response observed in *rhbdl2*−/− mutants. To test whether elevated Rac2 levels are required for this phenotype, we knocked down Rac2 in the *rhbdl2*−/− background. As expected, *rhbdl2*−/− + control MO exhibited a significant increase in wound-associated area compared with WT + control MO, while WT +Rac2 MO caused a decrease in wound-repaired area in comparison with WT+ control (**Fig. 6 D**). Strikingly, *rhbdl2*−/− + Rac2 MO significantly suppressed the enhanced wound repair phenotype, restoring wound area measurements to levels indistinguishable from WT (**Fig. 6 D**). Together, these results indicate that Rac2 contributes to enhanced wound repair in Rhbdl2-deficient larvae. Because our Rac2 knockdown approach was performed globally rather than in a macrophage-specific manner, we sought to directly test whether the enhanced wound repair phenotype in *rhbdl2*−/− larvae is macrophage dependent. To deplete macrophages, we injected mpeg:dsRed WT and *rhbdl2*−/− larvae with clodronate liposomes or PBS liposomes as a control at 2 dpf, screened for loss of macrophages at 24 h after injection, and performed tail transections at 3 dpf. Fluorescence imaging confirmed successful macrophage depletion at the wound site in clodronate-injected larvae at 72 hpw (**Fig. 6 E**), and quantification of mpeg: dsRed-positive macrophages confirmed significant reduction in both genotypes compared with PBS controls (**Fig. 6 F**). While clodronate-mediated macrophage depletion reduced wound repair area in both WT and *rhbdl2*−/− larvae, consistent with a general requirement for macrophages in regeneration, *rhbdl2*−/− larvae continued to display a slight but elevated wound repair area compared with clodronate-treated WT controls (**Fig. 6 G**). Collectively, these findings demonstrate that Rac2-dependent macrophage recruitment is a key driver of the enhanced wound repair response in *rhbdl2*−/− larvae, while the partial persistence of the phenotype following macrophage depletion points to additional macrophage-independent contributions, highlighting the multifaceted role of Rhbdl2 in regulating the cellular and molecular processes that govern wound repair.

**Figure 6.**
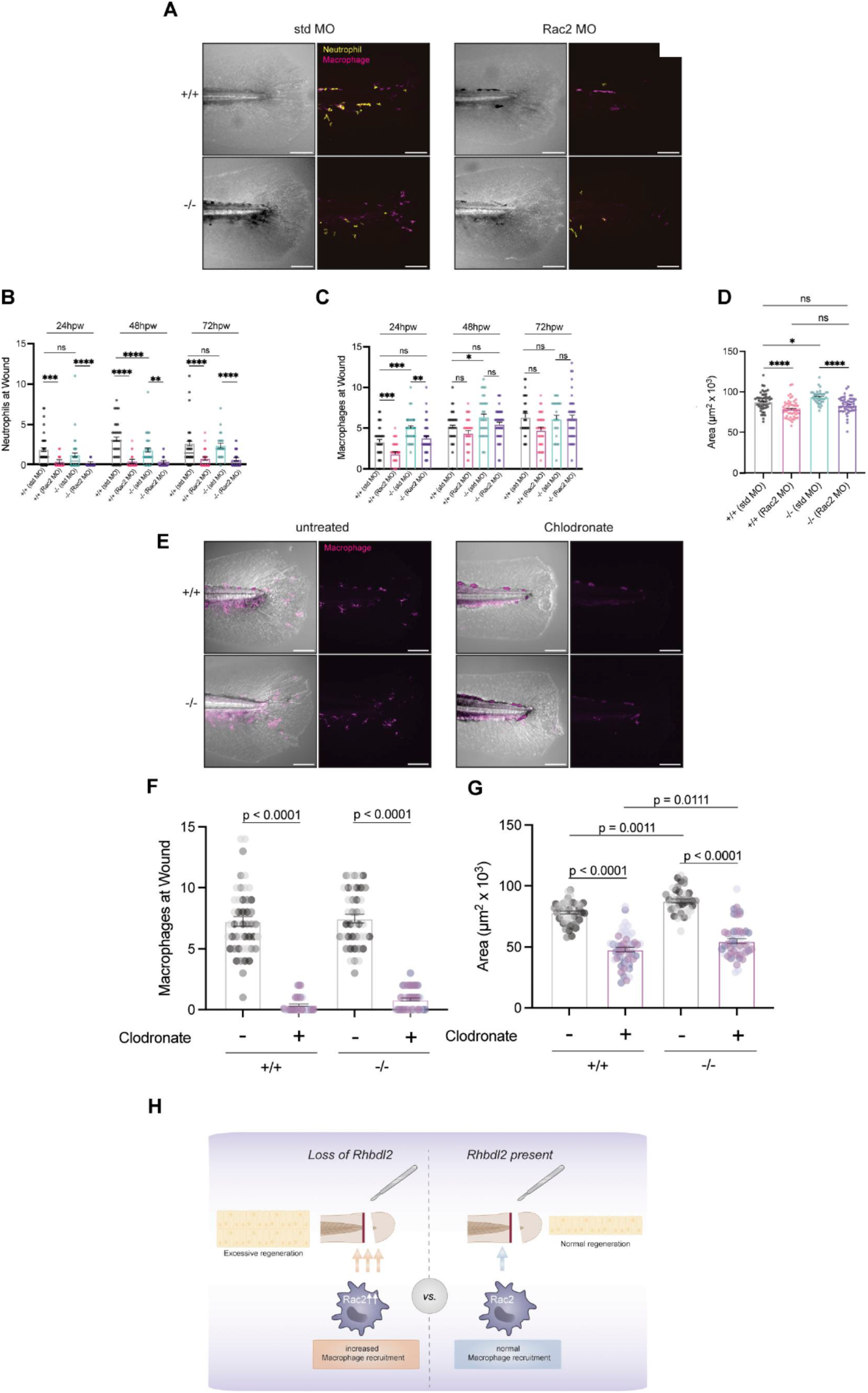
Rac2 ablation in *rhbdl2* mutants rescues wound repair and macrophage recruitment phenotype. (A) Representative images of WT or *rhbdl2*−/−in (Tg(lyz:GFP;mpeg1:dsRed)) background injected with either standard or Rac2 MO. Bright-field images (left) shown alongside merged neutrophil (yellow) and macrophage (magenta, right) images at 72 hpw. (B and C) Quantification of neutrophils (B) or macrophages (C) at wound sites in Rac2 MO–injected samples from 24 to 72 hpw. (D) Quantification of wound-repaired area in Rac2 MO–injected samples at 72 hpw. Data are from three independent experiments, (+/+: n = 51 [std MO], 56 [Rac2 MO]; −/−: n = 40 [std MO], 52 [Rac2 MO]). Lsmeans (±SEM) are reported, and P values calculated by ANOVA with Tukey’s multiple comparisons for all experiments. Scale bar = 100 µm. (E) Representative fluorescence images of mpeg:dsRed macrophages in WT and *rhbdl2*−/−larvae injected with PBS liposomes or clodronate liposomes at 72 hpw. Scale bar = 100 µm. (F) Quantification of macrophage number at the wound site at 72 hpw in WT and *rhbdl2*−/− larvae injected with PBS or clodronate liposomes. N = 3 independent replicates, n = 47 (WT PBS), n = 53 (WT clodronate), n = 41 (*rhbdl2*−/− PBS), and n = 47 (*rhbdl2*−/− clodronate). Lsmeans (±SEM) reported; P values calculated by ANOVA with Tukey’s multiple comparisons. (G) Quantification of wound repair area at 72 hpw in WT and *rhbdl2*−/− larvae injected with PBS or clodronate liposomes. N = 3 independent replicates, n = 47 (WT PBS), n = 53 (WT clodronate), n = 41 (*rhbdl2*−/− PBS), n = 47 (*rhbdl2*−/− clodronate). Lsmeans (±SEM) are reported; P values calculated by ANOVA with Tukey’s multiple comparisons. (H) Cartoon depicting the in vivo model of Rhbdl2-mediated wound repair. Injury is induced below the notochord in the tail fin. Loss of Rhbdl2 leads to elevated Rac2 protein levels and increased macrophage accumulation at the wound site, resulting in enhanced regenerative outgrowth. hpw, h post-wounding.

## Discussion

In this study, we identify the rhomboid intramembrane protease Rhbdl2 as a previously unrecognized modulator of wound-induced inflammation and wound repair and tail fin regeneration in zebrafish. Although Rhbdl2 is dispensable for normal development and fin growth, its loss is associated with enhanced tissue repair following injury, accompanied by heightened proliferation and apoptosis within the recovering tissue. Through unbiased proteomic profiling and in vivo live cell tracking and imaging, we uncover a central role for Rhbdl2 in shaping macrophage motility and recruitment during wound repair. Specifically, Rhbdl2-deficient larvae exhibit temporally dysregulated immune responses, marked by increased macrophage migration speed and recruitment at later stages, without corresponding changes in total leukocyte numbers. Mechanistically, our data show that Rac2 protein levels are elevated in *rhbdl2*−/− mutants and MO-mediated knockdown of Rac2 normalizes macrophage recruitment and suppresses enhanced wound repair (**Fig. 6 E**), consistent with Rac2 being required for the enhanced wound repair phenotype in *rhbdl2*−/− larvae. Together, these findings position Rhbdl2 as a critical modulator of immune cell behavior that constrains reparative growth. Previous studies have implicated Rhbdl2 in wound repair, but with conflicting results. Loss of Rhbdl2 function in HaCaT keratinocytes impaired scratch wound closure, suggesting a pro-repair role in epithelial cell migration (Cheng et al., 2011). In a contrasting study, suppressing the activity of human Rhbdl2 in HaCaT cells promoted cell migration by the activation of EGFR signaling, and Rhbdl2 deficiency impaired differentiation of human skin equivalents (Johnson et al., 2026). In congruence with this latter study, our in vivo analyses reveal that loss of Rhbdl2 in zebrafish is associated with enhanced wound repair following injury. This finding is unlikely to reflect genetic compensation or redundancy, as we generated a promoterless, RNA-null *rhbdl2* allele and confirmed the complete absence of *rhbdl2* mRNA, alongside validation with an Rhbdl2-targeted inhibitor. Moreover, expression levels of other rhomboid family members remained unchanged in Rhbdl2-deficient larvae, ruling out compensatory upregulation as an explanation for the phenotype. Instead, our findings highlight the importance of studying Rhbdl2 function within a multicellular, physiological context. Unlike in vitro scratch assays, which primarily capture cell-autonomous migratory behavior, the zebrafish wound repair model allows simultaneous interrogation of epithelial repair, immune cell dynamics, and tissue-scale reparative responses. Our data indicate that Rhbdl2 plays a critical role in coordinating leukocyte behavior, particularly macrophage recruitment, during wound repair, a layer of regulation that is absent in isolated cell culture systems. We show that gross accumulation of macrophages at the wound site is associated with enhanced tissue repair in Rhbdl2-deficient larvae. At first glance, this finding appears to contrast with prior studies in zebrafish demonstrating that persistent accumulation of pro-inflammatory (M1-like) macrophages impairs wound repair rather than enhancing it (Spencer et al., 2026). However, in *rhbdl2*−/− mutants, we observe that enhanced wound repair occurs in the context of heightened macrophage accumulation, suggesting that the reparative outcome is shaped not simply by macrophage persistent accumulation, but by the functional state of these cells. This interpretation is further supported by our live imaging data, which show that macrophages in *rhbdl2*−/− larvae exhibit a modest increase in migration speed toward the wound during the early recruitment phase (1–6 hpw), in addition to their increased accumulation and retention at the wound site at later time points (**Fig. 5, D and E**), consistent with an altered functional state rather than a simple increase in cell number. Consistent with this model, a recent study in zebrafish larvae reported that loss of the lipid-sensing receptor Gpr132b promotes a pro-resolving, pro-healing macrophage response characterized by increased macrophage accumulation coupled with reduced inflammatory signaling (Mercado Soto et al., 2025, Preprint). Notably, this immune profile phenocopies the enhanced wound repair response observed in Rhbdl2-deficient larvae. Accordingly, to directly assess whether loss of Rhbdl2 alters macrophage polarization, we quantified pro-inflammatory (M1; tnfa) and pro-resolving (M2; il10 and tgfb1a) markers by RT-qPCR in WT and *rhbdl2*−/− larvae under both unwounded and wounded conditions. Because Rhbdl2 sheds EGFR and can process EGF-family ligands (**Fig. S1 E**; Adrain et al., 2011), and EGFR signaling drives epithelial proliferation and migration during wound repair, we also measured the EGFR ligands hb-egfa and egf as a transcript-level readout of this substrate pathway rather than as macrophage polarization markers. None of these markers differed significantly between genotypes in either condition (**Fig. S5**), indicating that the enhanced macrophage accumulation in *rhbdl2*−/− mutants is not accompanied by a detectable transcriptional shift in bulk M1/M2 polarization or EGFR-ligand expression. We note that these measurements were performed on whole-tail RNA and therefore reflect bulk transcript abundance rather than macrophage-intrinsic expression, such that subtle or cell type–restricted changes in polarization may be masked at this resolution. Future profiling of isolated or FACS-sorted wound macrophages, or single-cell transcriptomics, will be needed to resolve macrophage activation states at higher resolution. Rhbdl2 is an intramembrane protease known to cleave membrane-embedded receptors and signaling proteins, thereby releasing soluble signaling fragments into intracellular, luminal, or extracellular compartments. Notably, macrophage-depleted *rhbdl2*−/− mutants still possessed a modest enhancement in regenerative outgrowth, suggesting that Rhbdl2 also regulates wound repair through macrophage-independent mechanisms. Such mechanisms can be derived from prior in vitro studies, which demonstrated that Rhbdl2 cleaves the CRAC channel subunit Orai1 to control its abundance, with Rhbdl2 loss resulting in aberrant calcium signaling (Grieve et al., 2021). Given emerging evidence that Orai1-dependent calcium signaling influences macrophage dynamics and functional states during wound repair, an important avenue for future investigation will be to determine how Orai1-mediated calcium signaling integrates with the Rhbdl2–Rac2 axis in regulating macrophage behavior and reparative outcomes. The additional Rhbdl2 substrate examined in this study, Spint1 (Johnson et al., 2017), is also a candidate key regulator of wound repair. In zebrafish, Spint1 knockdown leads to chronic inflammation and disrupted skin architecture (Mathias et al., 2007). Consistent with these defects, Spint1 mutants exhibit heightened leukocyte recruitment and increased proliferation of basal keratinocytes (Carney et al., 2007), phenotypes that closely mirror those observed in our study. EGFR signaling likewise plays a central role in regulating keratinocyte proliferation and migration during wound repair (Pastore et al., 2008; Repertinger et al., 2004), and human Rhbdl2 has been shown to suppress EGFR signaling by secreting and accumulating the decoy EGFR ectodomain in keratinocytes (Johnson et al., 2026). In addition, overexpressed Rhbdl2 was shown to cleave EGF and secrete its bioactive form that can activate EGFR signaling (Adrain et al., 2011), and a number of other substrates implicated in epithelial homeostasis, such as DDR1 or IL6R (Johnson et al., 2017), indicating that multiple substrates can contribute to the observed phenotypes in different tissue contexts. Despite a lack of conclusive information, loss of Rhbdl2 may permit enhanced proliferative signaling during wound repair since Rhbdl2 and its described substrates Spint1 and EGFR have both been implicated in promoting cell proliferation and migration across multiple cancer contexts (Chen et al., 2022; Gómez-Abenza et al., 2019; Johnson et al., 2026, Preprint; Normanno et al., 2006). How Rhbdl2 differentially regulates wound repair versus pathological growth programs mechanistically in vivo, and the extent to which macrophages contribute to these outcomes, remains an important question for future investigation. Rhbdl2 is expressed predominantly in the epidermis during development, as revealed by in situ hybridization and single-cell RNA-sequencing analyses. Accordingly, our current working model posits that during wound repair, Rhbdl2 functions primarily within epithelial cells to regulate Rac2 levels in macrophages in a cell-extrinsic manner. Prior studies demonstrate that macrophages are key drivers of Rac activation in the contexts of both wound repair and tumor invasion (Ramakrishnan et al., 2025). Rac2 activity is tightly regulated by its dynamic cycling between membrane-associated active and cytosolic inactive states, processes controlled by guanine nucleotide exchange factors, GTPase-activating proteins, and scaffolding proteins (Bos et al., 2007). We speculate that Rhbdl2-dependent intramembrane proteolysis of one or more such regulatory components in epithelial cells may generate downstream signals that alter Rac2 membrane residence time and stability, thereby elevating Rac2 protein levels in the absence of Rhbdl2. It is important to note, however, that our whole-body knockout model does not allow us to distinguish between epithelial cell-autonomous and macrophage-autonomous functions of Rhbdl2, and we cannot exclude the possibility that Rhbdl2 acts directly within macrophages to regulate Rac2 stability or signaling. Dissecting the cell type–specific contributions of Rhbdl2 and identifying its relevant substrates will be important directions for future work. Notably, attempts to reconstitute Rhbdl2 by injection of *rhbdl2* mRNA were precluded by dose-dependent developmental toxicity and lethality, with fluorescently tagged Rhbdl2 confirming that embryos with strong expression did not survive past 1 dpf (data not shown). This observation indicates that Rhbdl2 levels are tightly regulated during early development and that its overexpression is not tolerated. Cell type– specific, inducible transgenic strategies will therefore be required to achieve gain-of-function reconstitution in future work. Beyond its role in zebrafish wound repair, our findings suggest that Rhbdl2 functions as a negative regulator of macrophage accumulation during the resolution phase of wound repair, constraining macrophage-driven reparative programs. In pathological contexts, dysregulated macrophage recruitment and polarization are central drivers of chronic inflammation, fibrosis, and tumor progression. The ability of Rhbdl2 to restrain macrophage accumulation through a mechanism that involves Rac2 and promote appropriate resolution of the reparative response positions this protease as a potential molecular brake on maladaptive wound-healing programs (**Fig. 6 H**). Rhbdl2 therefore functions as a suppressor of reparative responses by restraining macrophage accumulation and limiting excessive tissue growth. Loss or inhibition of Rhbdl2 activity enhances macrophage persistence at wound sites and promotes wound repair outgrowth, suggesting that transient suppression of Rhbdl2 could be leveraged therapeutically to improve wound repair in contexts where repair is impaired. More broadly, our work identifies intramembrane proteolysis as an unexpected but powerful mechanism by which immune cell behavior is regulated, opening new avenues for therapeutic intervention in inflammatory, fibrotic, and cancer-associated pathologies.

**Figure S5.**
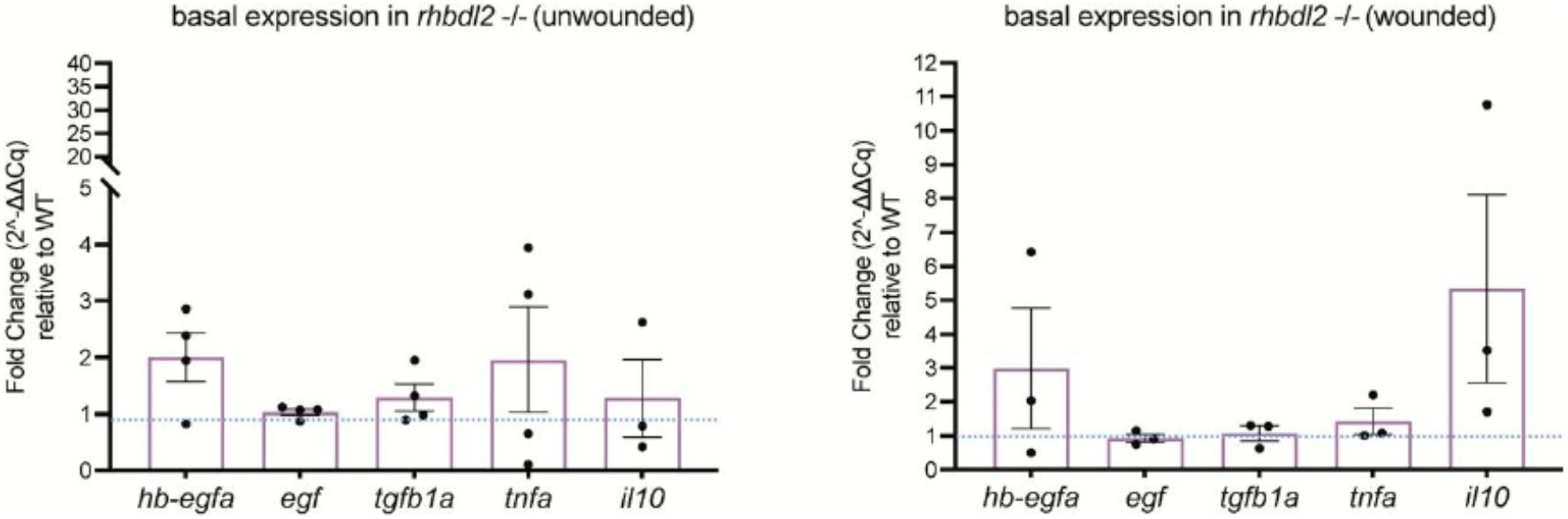
Macrophage polarization and EGFR ligand markers are unchanged in *rhbdl2*−/− mutants. RT-qPCR quantification of pro-inflammatory (M1; tnfa), pro-resolving (M2; il10, tgfb1a), and EGFR ligand (hb-egfa, egf) transcripts in *rhbdl2*−/− larvae relative to WT siblings, in unwounded (4 dpf) and wounded (24 hpw) conditions. Data are expressed as mean fold change relative to wild type (dashed line at 1; N = 3–4 independent biological replicates, n = 50 larvae per condition). Statistical significance was assessed by one-sample t test against a hypothetical mean of 1; no comparison reached significance (P > 0.05 for all genes in both conditions). Data are shown as mean ± SEM. hpw, h post-wounding.

## Materials and methods

### Zebrafish maintenance and handling

All zebrafish work follows protocols approved by the Institutional Animal Care and Use Committee at the University of California San Diego, San Diego, CA, USA (#S20060). Wild-type zebrafish AB lines were used, as well as the previously published transgenic lines Tg(lyz:EGFP), RRID:ZFIN_ZDB-ALT-071109-2 (Hall et al., 2007) and Tg(mpeg1:dsRed), RRID:ZFIN_ZDB-FISH-150901-6828 (Ellett et al., 2011). Adult fish were maintained on a 14-h/10-h light/dark schedule. Following breeding, fertilized embryos were maintained in E3 medium. To prevent pigment formation, larvae were maintained in E3 containing 0.2 mM N-phenylthiourea (PTU; Sigma-Aldrich) beginning at 24 h after fertilization. Prior to all experimental procedures, larvae were anesthetized in E3 water containing 0.2 mg/ml Tricaine (ethyl 3-aminobenzoate; Sigma-Aldrich).

### Zebrafish and human RHBDL2 alignment

Predicted transcripts and protein sequences of RHBDL2 are listed in Table 1. Transcript annotations are from GRCz11 for zebrafish, with their corresponding IDs from GRCz10 listed in **Fig. S1** when applicable. A MUSCLE alignment was generated in Jalview using the protein sequences of mature human RHBDL2 (UniProtKB accession no. Q9NX52; residues 1–303) and mature zebrafish RHBDL2 (UniProtKB accession no. F1QAX9; residues 1– 293; UniProt Consortium 2017). The 191 identical residues of human and zebrafish RHBDL2 (65% sequence identity) were mapped onto the predicted AlphaFold model of human RHBDL2 (Tunyasuvunakool et al., 2021).

**Table 1.**
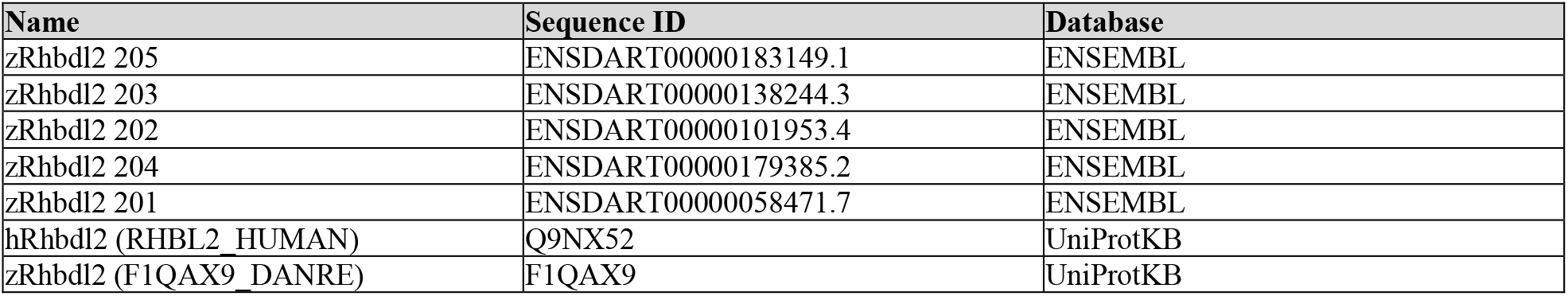
Annotated RHBDL2 sequences used in this study.

### Tissue culture and western blotting

HEK293T cells (CRL-3216; ATCC) were incubated in DMEM supplemented with 10% FBS at 37°C and 5% (vol/vol) CO2 in a humidified incubator. All genes were cloned into the pcDNA3.1 vector as described (Johnson et al., 2017) and sequence verified. One day before transfection, HEK293T cells were seeded into a 6-well plate at a density of 5 × 105 cells per well. The transfection reaction was prepared in 100 μl serum-free DMEM (SFM) with 3 μl of FuGENE 6, 600 ng of Spint1 expression plasmid, and 300 ng of either human or danio RHBDL2 expression plasmids, respectively. One day after transfection, cells were washed with PBS and incubated in 750 μl of SFM for another 24 h. The conditioned SFM was harvested, centrifuged at 2,000 × g to remove residual detached cells and debris, and proteins were precipitated with trichloroacetic acid as described (Johnson et al., 2017). Cell lysates were harvested by lysing the attached cell monolayers in 150 μl SDS PAGE sample buffer per well. Protein samples were resolved by SDS-PAGE alongside a molecular weight protein ladder, then transferred to PVDF membranes. Membranes were blocked in 5% nonfat milk in TBST for 1 h at RT, incubated with primary antibodies overnight at 4°C, washed three times in TBST, and incubated with HRP-conjugated secondary antibodies for 1 h at RT. Blots were developed by chemiluminescence and imaged on a ChemiDoc MP (Bio-Rad). HaCaT cells (300493; Cell Lines Service GmbH) deficient in *rhbdl2* and their derivative stably expressing zebrafish Rhbdl2 were maintained in DMEM supplemented with 10% FBS at 37°C and 5% (vol/vol) CO2. For EGFR cleavage assays, HaCaT cells were seeded into 6-well plates at a density of 1 × 106 cells per well. The next day, cells were washed with PBS and incubated in 750 μl SFM with or without STS1650 (50 µM) for 24 h. Conditioned media were harvested and processed as described above. Cell lysates and conditioned media were analyzed by Western blotting using antibodies against endogenous EGFR (ab264540; Abcam, mouse, RRID: AB_3665828) and HA (Rhbdl2) (3724; Cell Signaling Technology, rabbit, RRID: AB_1549585) (Johnson et al., 2017).

### Generation of *rhbdl2* mutant line and genotyping

We generated a stable promoterless *rhbdl2* mutant allele, designated sd73. Zebrafish mutants were generated using CRISPR/Cas9 single-cell injections. gRNAs for zebrafish Rhbdl2 (ENSDART00000183149.1) were designed using CRISPRscan (Moreno-Mateos et al., 2015) to target 5′ upstream of exon 1 and within exon 2. The 5′ target sequence was 5′-AGCTTGCATCGT AAAGGCTG-3′ while the exon 2 target sequence was 5′-GGAGTT ACATGAGACTTATC-3′. The gRNAs and Cas9 protein (Integrated DNA Technologies) were prepared by the manufacturer’s instructions and combined to a final RNP concentration of 1.5 µM. The RNP solution was injected into one-cell AB embryos in a 1 nl volume. To confirm genome editing, injected 2 dpf embryos had genomic DNA extracted and amplified using the primers listed below before being sent out for Sanger sequencing: Rhbdl2 F: 5′-GTGACACTGGAGCAGCCTAATG-3′, and Rhbdl2 R: 5′-CTTTTC AGCTTTCTTTCCAGCCTTC-3′. F0 mosaic cuts were confirmed with gel electrophoresis and sequencing, with clutches having confirmed CRISPR cuts grown to adulthood. Adult F0 CRISPR-injected fish were screened for germline mutation by outcrossing and testing individual larvae using the primers above and sequencing. Heterozygous *rhbdl2*+/− fish were maintained by outcrossing F1 CRISPR mutants to AB WT background zebrafish before being genotyped by genomic DNA extraction from fin clips. PCR was carried out using the primers listed above along with an additional set to detect deletion of the genomic region of interest: R2 Mutant F: 5′-GAA TTAAGTTACCCGTTTATGTG-3′, and R2 Mutant R: 5′-CTTTGC AGAAGGACGAAGAAG-3′. PCR products were separated on a 1.5% agarose gel for 30 min to determine individual fish genotypes. For experimental purposes, F2 or F3 heterozygotes were incrossed to produce a generation of adult homozygous WT and mutant siblings. These adults were incrossed to produce WT and *rhbdl2*−/− clutches. These cousin larvae were only used for experimentation, while outcrossing of heterozygotes continued to be used to maintain subsequent generations.

### RT-qPCR

RNA and DNA were extracted from pooled samples of 10–20 larvae from homozygous incrosses using TRIZOL (Invitrogen) and following the manufacturer’s protocol. DNA underwent PCR amplification using Q5 High-Fidelity 2× Master Mix (New England Biolabs) as described above to confirm genotypes. Extracted RNA was used for cDNA production using High-Capacity cDNA Reverse Transcription Kit (Applied Biosciences). qPCR was performed using AzuraView GreenFast qPCR Blue Mix LR (Azura Genomics) and an OpusFlex384 (Bio-Rad). Primers for target genes are listed below. Data were normalized to rps11 using the ΔΔCt method to illustrate fold change over WT samples. For *rhbdl2* knockout validation, RNA was extracted from larvae at 2 dpf and 6 dpf. For wound-induced expression analysis of *rhbdl2*, incrosses of homozygous adult WT and *rhbdl2*−/− siblings were done to produce homozygous cousins of WT and mutants. Larvae at 2 dpf were injured with a transection wound at the tip of the notochord. At 24 hpw, fins were collected by transection of repairing tissue at the tip of the blood circulatory loop. 30–50 fins were pooled and flash-frozen, with unwounded 3 dpf control tissue collected at the same stage. RNA was extracted from fin tissue as described above. Primers for *rhbdl2* and rps11 are listed below, and data were normalized to rps11 and represented as fold change over unwounded WT, age-matched control tissue. RT-qPCR primers were used as followed: Rhbdl2 exon2-3 qF: 5′-TTCATCATCCTCATCAGTTT-3′, Rhbdl2 exon2-3 qR: 5′-GTT GTTCTGGTCTATAGGTA-3′; Rhbdl2 exon4 qF: 5′-CATAAAGGC TTTGAAGTTGG-3′, Rhbdl2 exon4 qR: 5′-CATAAGGGCATAAAC ACCAC-3′; Rhbdl1 exon2 qF: 5′-GTACAGATCATGGTGTTCAT-3′, Rhbdl1 exon2 qR: 5′-GCAGCACCCATTTATTGAGC-3′; Rhbdl3 exon1 qF: 5′-GTAGCTCCAGAGGACCAATG-3′, Rhbdl3 exon1 qR: 5′-CAGTTCTGATCCATGAGCTG-3′; Rhbdl4 exon4 qF: 5′-ACC CTGGGACATCATTTGTT-3′, Rhbdl4 exon4 qR: 5′-CCCTCCATA ATGCCCATTAG-3′; rac1a qF: 5′-TCCGGCCTCATTTGAAAACG-3′, rac1a qR: 5′-GTGTCCTTGTCATCTCGCAA-3′; rac1b qF: 5′-CTC TCTCGTACCCTCAGACG-3′, rac1b qR: 5′-TGGGAGTGGTTTGAC AGTGA-3′; rac2 qF: 5′-AATACCTGGAGTGTTCGGCC-3′, rac2 qR: 5′-GCCCTTCTTCTTGACCTTGG-3′; rac3a qF: 5′-CCCAGGAGA GTATATACCCACAG-3′, rac3a qR: 5′-GGTATGAAAGTGGTCTAA GGCG-3′; rac3b qF: 5′-CCCCGGCGAGTACATCCCTACAG-3′, rac3b qR: 5′-GGTATGAGAGTGGACGGAGGCG-3′; hb-egfa qF: 5’GGGCCCTCATGCATATGTGA-3′, hb-egfa qR: 5′-AGAATTTCC ACACGGTCGCC-3′ (Cigliola et al., 2023); egf qF: 5′-TAAGTG AGTGGACAATGTT-3′, egf qR: 5′-GTCTTCGTGTTCCATCTA-3′ (Sun et al., 2018); tgfb1a qF: 5′-TACGTGGGAAGGCAACAC AA-3′, tgfb1a qR: 5′-CCCGACTGAGAAATCGAGCC-3′(Tian 田□波 et al., 2026); tnfa qF: 5′-GCGCTTTTCTGAATCCTA CG-3′, tnfa qR: 5′-TGCCCAGTCTGTCTCCTTCT-3′ (Candel et al., 2014); il10 qF: 5′-ATTTGTGGAGGGCTTTCCTT-3′, il10 qR: 5′-AGAGCTGTTGGCAGAATGGT-3′ (Galindo-Villegas et al., 2012); rps11 qF: 5′-TAAGAAATGCCCCTTCACTG-3′, rps11 qR: 5′-GTCTCTTCTCAAAACGGTTG-3′(de Oliveira et al., 2015).

### Developmental imaging

Homozygous cousin larvae were generated from adult WT and *rhbdl2*−/− siblings as described above. Larvae were raised to 2 dpf without PTU and anesthetized before imaging on a Zeiss Discovery. V8 stereoscope with an Apo S 1.0× FWD 60 mm objective (Zeiss). Images were taken at 3.2× magnification using an Axiocam 208 color camera (Zeiss) and Labscope version 4.2.1 (Zeiss). Imaging was performed at RT (~22°C). Adult fish were raised to 6 mpf and separated by sex. Adult fish were anesthetized in Tricaine (0.017% in E3 media) and placed on petri dish with white tape underneath for best visualization. Water was removed to leave a small amount to cover the fish and allow for respiration. Fish were briefly imaged using an iPhone 13 camera with a ruler placed adjacent for size measurements.

### Wound repair assays

For larval wound repair assays, incrosses of adult F3 WT and *rhbdl2*−/− siblings were set up to produce homozygous cousins possessing WT or promoterless alleles. Larvae were dechorionated at 2 dpf and transferred to petri dishes previously coated in milk. Tail transections were performed at 3 dpf using a surgical blade (feather no.:10) to create a vertical cut at the posterior tip of the developing notochord in the caudal fin without injuring the notochord. Larvae were washed in E3 and allowed to repair for up to 4 dpw. At indicated times, larvae were fixed with 4% PFA (Sigma-Aldrich) in PBS overnight at 4°C, followed by washing three times in PBS. Larvae were prepared for imaging by bisecting larvae at the posterior tip of the yolk sac in order for fins to lie flat before transferring to a 35-mm glass-bottom imaging dish (Willco). Fins were imaged in PBS at RT with an AX-R point-scanning confocal on a Ti2-E inverted microscope (Nikon). Images were acquired using a 20× Plan Apo Lambda D objective (Nikon, NA=.080) as 50 µm z-stacks with 5-µm optical sections and 512 × 512 resolution. Images were acquired and processed using NIS Elements Ar version 6.10.01. Unwounded age-matched larvae at 6 dpf were measured as a developmental control.

### Rhbdl2 ketoamide inhibition wound assay

WT AB larvae were raised to 2 dpf and transferred to 24-well plates in groups of ~10 larvae per well. Larvae were treated with STS1650 (50 µM) or an equivalent volume of DMSO in E3-PTU medium beginning at 2 dpf. *rhbdl2*−/− larvae treated with DMSO served as a positive control. Wounding was performed at 3 dpf using a surgical blade to create a vertical transection at the tip of the notochord to standardize wound location. Larvae were allowed to regenerate for 72 hpw, after which they were fixed in 4% PFA (Sigma-Aldrich) in PBS overnight at 4°C, then washed three times in PBS. The wound repair area was quantified as described above.

### Caspase inhibition wound assay

WT AB and *rhbdl2*−/− mpeg:dsRed larvae were generated and wounded at 3 dpf as previously described. Following wounding, larvae were transferred to solutions of the pan-caspase inhibitor QVD-OPH (50 µM) or an equivalent volume of DMSO in E3-PTU medium. Larvae were fixed at 72 hpw in 4% PFA (Sigma-Aldrich) in PBS overnight at 4°C, then washed three times in PBS. The wound repair area was quantified as described above. Macrophage number at the wound site was determined by counting mpeg:dsRed-positive cells in the wound region using FIJI software.

### Clodronate liposome macrophage depletion wound assay

WT and *rhbdl2*−/− mpeg:dsRed larvae were injected in the caudal vein with clodronate liposomes or PBS liposomes at 2 dpf. At 3 dpf, larvae were screened for successful macrophage depletion by loss of mpeg:dsRed fluorescence prior to wounding. Larvae were wounded at 3 dpf as previously described and allowed to regenerate for 72 hpw. Larvae were fixed in 4% PFA (Sigma-Aldrich) in PBS overnight at 4°C, then washed three times in PBS. The wound repair area was quantified as described above. Macrophage number at the wound site was determined by counting mpeg:dsRed-positive cells in the wound region using FIJI software.

### EdU labeling and immunohistochemistry

Adult F3 WT and *rhbdl2*−/− siblings were incrossed to produce homozygous cousin offspring. Proliferation in the fin was measured with the Click-iT Plus EdU Imaging Kit (Life Technologies). After transection wounding as described above, wounded larvae were incubated in a 250 µM EdU solution in E3 media for 6 h at RT. Wounded larvae were incubated from 66 to 72 hpw alongside age-matched unwounded controls. Larvae were fixed overnight in 4% PFA in PBS at 4°C before storage in methanol at −20°C until staining. Larvae were rehydrated in a series of 5-min washes at ratios of 2:1, 1:1, and 1:2 methanol: 0.2% Triton X-100 in PBS (PBSTx) followed by 30-min washes twice in PBSTx. Larvae were blocked in a 1.5% sheep serum, 1% BSA, PBSTx for 3 h before being stained according to the manufacturer’s instructions overnight at 4°C in the dark. All incubations included slight agitation by rocking. Transection of tails at the yolk sac and imaging in a glass-bottom dish were done as described above. Immunofluorescence images were acquired using an AX-R point-scanning confocal on a Ti2-E inverted microscope (Nikon) equipped with an LUA-S4 laser unit with 405/ 488/561/640 lines. Images were acquired with 20× air objective (CFI Plan Apo Lambda D, numerical aperture of 0.8; Alexa Fluor 647 picolyl azide fluorophore used for EdU detection) as 50-µm z-stacks, 3-µm optical sections, and 512 × 512 resolution.

### Mitosis and apoptosis labeling

Homozygous cousin larvae were generated and wounded at 3 dpf as previously described. Larvae were fixed at either 24 or 72 hpw in 4% PFA in PBS overnight at 4°C before transfer to methanol at −20°C. Larvae were rehydrated in a series of 5-min washes at ratios of 2:1, 1:1, and 1:2 methanol: 0.3% Triton X-100, 1% DMSO, 0.05% Tween-20 in PBS (PDT solution) followed by washing twice in PDT for 30 min at RT. Larvae were blocked in a solution of 10% heat-inactivated FBS, 2% BSA, and 0.05% Tween-20 in PBS for 1 h at RT. Samples were co-stained with monoclonal rabbit anti-active Caspase3 antibody (#559565; BD Biosciences, RRID: N/A, product discontinued) diluted 1:500 and monoclonal rat anti-phospho-H3 (pSer 28) antibody (H9908; Sigma-Aldrich, RRID:AB_260096) diluted 1:300, in blocking overnight at 4°C. Samples were washed twice in PDT for 30 min at RT and incubated in blocking again for 1 h. Incubation with DyLight donkey anti-rabbit 550 (Thermo Fisher, SA510039, RRID: AB_2556619) and DyLight donkey anti-rat

488 secondary antibodies (SA5-10038; Thermo Fisher Scientific, RRID: AB_2556618) diluted 1: 300 was performed overnight at 4°C. Samples were washed three times in PDT before imaging in a glass-bottom dish using the laser scanning microscope with Plan-Apochromat NA 0.8/ 20× air objective, as described above.

### Live Rhobo6 staining and imaging

Homozygous cousin larvae were generated and wounded at 3 dpf as previously described. Wounded larvae were grown to 72 hpw whereby they were anesthetized and loaded onto a plate of 3% agarose in E3/MB with a small amount of water. Rhobo6 dye was prepared and diluted to 1 mM in DMSO. A final solution at 100 µM was made with a 1:9 mixture of dye with phenol red, and 3 nl was injected into the caudal vein of the 72 hpw larvae. Confirmation of injection was detected by the presence of phenol red in the heart of larvae. Larvae were allowed to recover for 30 min in E3/MB supplemented with Rhobo6 (5 µM). Larvae were mounted on their side in 3% low-melting point agarose with tails open by manually removing agarose around the area. Dishes were covered in E3 and mounted 10–20 min before performing two-photon imaging on a Thorlabs Bergamo II multiphoton microscope and a fs-pulsed 80-Mhz Ti:S laser (MaiTai DeepSee; Spectra-Physics). Live images were captured using an excitation wavelength of 1,080 nm (<10 mW of excitation power) and a 16×/0.8 W Nikon water immersion objective (CF175 LWD). Imaging was controlled by ThorImage 4.3. Emitted light was collected using Thorlabs PMT2100 dual-channel GaAsP photomultiplier tubes (PMTs) at a controlled RT of 28.5°C. Frames were 1,024 × 512 pixels covering a 454.75 × 232.38 µm field of view (1.8× magnification) over 50–80 µm in Z (resulting in a total imaged volume of 5.28 × 106 to 8.45 × 106 μm3), collected at a rate of 30 Hz, and averaged 40 times to increase signal to noise. Images were subsequently processed in ImageJ to produce average z-stacks over the entire Z-stack and then blinded and scored in ImageJ.

### Mass spectroscopy

#### Sample preparations

Samples were desiccated and reconstituted in 200 µl of 6 M guanidine-HCl. The samples were then boiled for 10 min, followed by 5 min of cooling at RT. The boiling and cooling cycle was repeated a total of 3 cycles. The proteins were precipitated with the addition of methanol to a final volume of 90%, followed by vortexing and centrifugation at maximum speed on a benchtop microfuge (14,000 rpm) for 10 min. The soluble fraction was removed by flipping the tube onto an absorbent surface and tapping to remove any liquid. The pellet was suspended in 200 µl of 8 M Urea made in 100 mM Tris pH 8.0. TCEP was added to a final concentration of 10 mM, and chloroacetamide solution was added to a final concentration of 40 mM, and vortexed for 5 min Three volumes of 50 mM Tris, pH 8.0, were added to the sample to reduce the final urea concentration to 2 M. Trypsin was in 1:50 ratio and incubated at 37°C for 12 h. The solution was then acidified using TFA (0.5% TFA final concentration) and mixed. The sample was desalted using C18-StageTips (Thermo Fisher Scientific) as described in the manufacturer’s protocol. The peptide concentration of the sample was measured using BCA.

#### Mass spec acquisition

DDA; 1 µg of each sample was analyzed by ultrahigh-pressure liquid chromatography (UPLC) coupled with tandem mass spectrometry (LC-MS/MS) using nanospray ionization. The nanospray ionization experiments were performed using a timsTOF 2 HT hybrid mass spectrometer (Bruker) interfaced with nanoscale reversed-phase UPLC (EVOSEP ONE). Evosep method 15 samples per day was utilized using a 15 cm × 150-μm reversed-phase column packed with 1.5-μm C18 beads (PepSep, Bruker) at 58°C. The analytical columns were connected to a fused silica ID emitter (10 μm ID; Bruker Daltonics) inside a nanoelectrospray ion source (Captive spray source; Bruker). The mobile phases comprised 0.1% FA as solution A and 0.1% FA/ 99.9% ACN as solution B. The mass spectrometry settings for the TimsTOF Pro 2 are as follows: PASEF method for standard proteomics. The values for mobility-dependent collision energy ramping were set to 95 eV at an inverse reduced mobility (1/k0) of 1.6 V s/cm2 and 23 eV at 0.73 V s/cm2. Collision energies were linearly interpolated between these two 1/k0 values and kept constant above or below. No merging of TIMS scans was performed. Target intensity per individual PASEF precursor was set to 20,000. The scan range was set between 0.6 and 1.6 V s/cm2 with a ramp time of 166 ms. 14 PASEF MS/MS scans were triggered per cycle (2.57 s) with a maximum of seven precursors per mobilogram. Precursor ions in an m/z range between 100 and 1,700 with charge states ≥3+ and ≤8+ were selected for fragmentation. Active exclusion was enabled for 0.4 min (mass width 0.015 Th, 1/k0 width 0.015 V s/cm2). Peptide identification was carried out using Peaks Studio 12.5 (Bioinformatics Solutions Inc.).

#### Mass spectrometry analysis

To evaluate proteomic changes, adult F3 WT and *rhbdl2*−/− siblings were incrossed to produce homozygous cousin offspring. Larvae were raised without PTU until 2 dpf before anesthetization. 50–60 larvae at 2 dpf were pooled and deyolked in Ringer’s solution with gentle disruption using a P1000 pipette tip. Larvae were washed twice in PBS, and the solution was removed before samples were stored at −80°C until submission for LC-MS/MS analysis. DB values for each protein were generated for three replicates each of both WT and *rhbdl2*−/− samples. These estimated abundance measures were merged across all samples and used as input to the Perseus software package (Tyanova et al., 2016) for processing. Abundance values were log-transformed, and missing values for proteins in samples where the protein was undetected were imputed. Imputed values were assigned using the Perseus default method, which randomly assigns values from a Gaussian distribution with a −0.3 sd downshift. Lastly, the values were normalized using a “width adjustment” before being exported to the R statistical environment (www.cran.org). Within R, the data were used in differential abundance analysis using the limma package (Ritchie et al., 2015) to compare the WT and *rhbdl2*−/− groups. Significantly differentially abundant proteins (P-value <0.05) were used as input for enrichment analysis in the GO, KEGG, and Reactome ontologies with the g:Profiler tool (https://biit.cs.ut.ee/gprofiler/gost, [Peterson et al., 2020]). Scatter plots and box plots were made with the ggplot2 package in R, while the heatmap was made with the gplots package.

#### Leukocyte imaging

For analysis of leukocytes, a transgenic line was developed by crossing *rhbdl2*+/− to AB WT zebrafish labeled with fluorescent neutrophils (Tg(lyz:GFP)) or fluorescent neutrophils and macrophages (Tg(lyz:GFP;mpeg1:dsRed)). Genotyped heterozygotes were incrossed to produce homozygous, fluorescently labeled adults that were incrossed again to produce WT and *rhbdl2*−/− cousins for experimental use. Wounding and fixing were performed as described above before fins were imaged on the laser scanning confocal as described for wound repair assays. Whole larvae were live imaged in a zWEDGI (Huemer et al., 2017) at indicated stages and acquired on the point-scanning confocal with a 10× plan Apo Lambda D air objective (180-µm z-stacks, 5-µm optical sections, 3 × 1 tiles with 15% overlap, 512 × 512 resolution). For time-lapse imaging of leukocyte recruitment, 3 dpf WT and *rhbdl2*−/− larvae were wounded by tail transection as described above. Larvae were mounted in zWEDGI devices (held in place using 1% low-melting-point agarose [Sigma-Aldrich] at the head). Images of the caudal fin were acquired at RT using the point-scanning confocal microscope (20× Plan Apo Lambda D air objective, 50-µm z-stacks, 5-µm optical sections, one z-stack every 2 min for 6 h at indicated times following injury, 512 × 340 resolution).

#### Morpholino injections

Previously published morpholinos were obtained from GeneTools for rac2, 5′-CCACCACACACTTTATTGCTTGCAT-3′ along with the standard control MO (GeneTools). Morpholinos were resuspended in water to a stock concentration of 1 mM. Morpholinos were further diluted in water with 0.5× CutSmart Buffer (New England Biosciences) and 0.1% phenol red to an injected concentration of 0.1 mM. Alongside the same concentration of control MO, 1 nl of diluted morpholino was injected at the 1-cell stage of fertilized embryos.

#### Image analysis and processing

All images were acquired at RT and processed using NIS Elements Ar version 6.10.01. Images are always shown with anterior to the left and dorsal at the top except where noted. Brightness and contrast were adjusted for best visualization and were not used during fluorescent analysis.

#### Wound repair area quantification

Wound repaired area was measured from the posterior tip of the notochord to the caudal edge of the tail fin with the polygon tool in FIJI image analysis software (Schindelin et al., 2012).

#### Immunohistochemistry

For EdU analysis, images were reconstructed in 3D using Imaris software (Bitplane, version 11.0). The Spots function was used to quantify the number of EdU-positive cells in the fin region distal to the blood circulatory loop, where xy = 7 µm and z = 14 µm. pH_2_-positive cells were counted manually using max projected images in the same area beyond the blood loop. Apoptosis activation was quantified on max projected images in FIJI by outlining the fin past the blood circulatory loop using the polygon tool in the corresponding bright-field image. Total threshold area for active-Casp3 signal was determined using the threshold plugin in FIJI by auto-thresholding with the “Yen Dark” method (Yen et al., 1995).

#### Leukocyte counts

Leukocyte numbers in wound assays were determined by hand in the region beyond the caudal vein tip using FIJI software on maximum intensity projections.

Total neutrophil and macrophage numbers were determined using Imaris (Bitplane, version 11.0) with the Spots function defined by xy = 10 µm and z = 20 µm. Leukocytes within the eye and heart were excluded. Full-size measurements were also taken on max projected images for 2 dpf length. Using time-lapse confocal images, leukocytes were tracked using the Spots function. The threshold for spot size was set at xy = 10 for neutrophils and xy = 8 for macrophages. The cells recruited to the wound area distal to the caudal vein loop were tracked manually.

#### Statistical analyses

All graphs were plotted on GraphPad (Prism v. 10). For all statistical analyses, at least three independent replicates were conducted, with a replicate defined as a separate clutch of larvae spawned on different days. RT-qPCR results were determined by comparing the calculated ΔCq values of the experimental conditions using a one-sample t test. Fold change was calculated and plotted in terms of mean with a 95% confidence interval. Differential protein abundances were compared using the limma package within R (Ritchie et al., 2015). For all other quantitative experiments, data were pooled from independent replicates and summarized in terms of least-squares adjusted means (emeans) and SEM (Golenberg et al., 2020; Vincent et al., 2016). These results were analyzed using ANOVA with Tukey’s multiple comparisons test. Graphical representation depicts individual data points color-coded to reflect replicates. Averages per replicate represented by solid color points superimposed on graphs. Statistical analysis and graphical representations were done using R version 4.5.2 and GraphPad Prism version 10. All t tests and ANOVA comparisons were reported in this study as two-sided, and data distribution was assumed to be normal, but this was not formally tested.

## Online supplemental material

This article includes five supplemental figures and two videos. **Fig. S1** shows characterization of zebrafish Rhbdl2 structure and function. **Fig. S2** shows characterization of the ECM matrix in wound-repaired fins of *rhbdl2*−/− mutants. **Fig. S3** shows differential abundance of proteins between WT and *rhbdl2*−/− larvae. **Fig. S4** shows development of leukocytes is unaffected in promoterless *rhbdl2*−/− mutants. **Fig. S5** shows macrophage polarization and EGFR ligand markers are unchanged in *rhbdl2*−/− mutants. Video 1 shows time-lapse imaging of leukocyte recruitment to the tailfin wound in a WT larva. Video 2 shows time-lapse imaging of leukocyte recruitment to the tailfin wound in a *rhbdl2*−/− larva.

## Data availability

The proteomics datasets generated during this study are available in the Mendeley Data repository at https://doi.org/10.17632/gwh3j3rrjr.1.

## Acknowledgments

We thank Dr. Majid Ghassemian of the Biomolecular and Proteomics Mass Spectrometry Facility for processing our samples for mass spectrometry. We also thank the Neal lab members for in-depth discussions and technical assistance and Dr. Kayvon Pedram, University of California San Diego, La Jolla, CA, USA for the Rhobo6 dye. We thank Dr. Cressida Madigan and members of the Madigan laboratory, including Dr. Sumedha Ravishankar and Dr. Megan Hayes, for mentorship and technical training provided to S.G. during the early phase of his graduate training, and for access to laboratory resources, including Imaris imaging and analysis software, used in this study. We also thank S.G.’s thesis committee members, Drs. Deborah Yelon, Jill Wildonger, and David Traver, for their guidance in the development of the promoterless *rhbdl2* allele and for their continued scientific input and project guidance over the course of this work.

These studies were supported by Pew Biomedical Award, Howard Hughes Medical Institute Freeman Hrabowski Program, National Institutes of Health (NIH) grant 1R35GM133565, Chan Zuckerberg Initiative grant, and National Science Foundation (NSF CAREER grant 2047391 (to S.E. Neal); Czech

Science Foundation (project 21-24456S) and institutional research concept RVO 61388963 to the Institute of Organic Chemistry and Biochemistry to K. Strisovsky; Packard Foundation Fellowship, Pew Biomedical Scholar Award, McKnight Scholars Award, the NIH New Innovator Award DP2EY036251, and the Kavli Institute for Brain and Mind to M. Lovett-Barron; and Postdoctoral Fellow of the Helen Hay Whitney Foundation to K. Martin.

## Author contributions

Saroj Gourkanti: conceptualization, data curation, formal analysis, investigation, methodology, project administration, resources, software, supervision, validation, visualization, and writing—original draft, review, and editing. Gayathri Ramakrishnan: data curation, formal analysis, investigation, methodology, and visualization. Jacqueline Cheung: formal analysis, investigation, methodology, resources, and writing—review and editing. Taylor J. Schoen: Investigation, Writing - review and editing. Yazmin Munoz: methodology and resources. Rosa M. Chavez: investigation. Jan Dohnálek: investigation and writing—review and editing. Katie Martin: data curation and investigation. Matthew Lovett-Barron: investigation, resources, and supervision. Thomas Whisenant: formal analysis, software, and writing— review and editing. Kvido Strisovsky: conceptualization, data curation, funding acquisition, methodology, project administration, resources, supervision, validation, and writing—original draft, review, and editing. Sonya E. Neal: conceptualization, data curation, funding acquisition, investigation, project administration, resources, supervision, and writing—original draft, review, and editing.

## Disclosures

All authors have completed and submitted the ICMJE Form for Disclosure of Potential Conflicts of Interest. K. Strisovsky reported a patent to US10927146B2 issued “Kvido Strisovsky” and a patent to EP3638686 (B1) issued Kvido Strisovsky. No other disclosures were reported.

## Notes

https://doi.org/10.17632/gwh3j3rrjr.2

### Competing Interest Statement

The authors have declared no competing interest.

### Summary of Updates

Figure 2 revised to include a new pharmacological inhibition experiment (STS1650) showing dose-dependent recapitulation of the rhbdl2 mutant enhanced wound repair phenotype in WT larvae. Figure 6 revised to include a clodronate liposome macrophage depletion experiment testing whether the enhanced wound repair phenotype is macrophage-dependent. Figure S1 updated to include validation that STS1650 inhibits zebrafish Rhbdl2 proteolytic activity (EGFR ectodomain shedding assay). Figure S3 updated to include a pan-caspase inhibitor (QVD-OPH) experiment testing whether apoptosis drives the increased macrophage accumulation phenotype. Results section updated to describe all three new drug-treatment experiments and their findings. Materials and methods updated to add protocols for STS1650 treatment, clodronate liposome macrophage depletion, and caspase inhibitor (QVD-OPH) treatment.

